# Brain neuromarkers predict self- and other-related mentalizing across adult, clinical, and developmental samples

**DOI:** 10.1101/2025.03.10.642438

**Authors:** Dorukhan Açıl, Jessica R. Andrews-Hanna, Marina Lopez-Sola, Mariët van Buuren, Lydia Krabbendam, Liwen Zhang, Lisette van der Meer, Paola Fuentes-Claramonte, Edith Pomarol-Clotet, Raymond Salvador, Martin Debbané, Pascal Vrticka, Patrik Vuilleumier, David A. Sbarra, Andrea M. Coppola, Anita Tusche, Lars O. White, Tor D. Wager, Leonie Koban

## Abstract

Human social interactions rely on the ability to reflect on one’s own and others’ internal states and traits—a process known as mentalizing. Impaired or altered mentalizing is a hallmark of multiple psychiatric and neurodevelopmental conditions. Yet, replicable and easily testable brain markers of mentalizing have so far been lacking. Here, we apply an interpretable machine learning approach to multiple datasets (total *N*=390) to train and validate fMRI brain signatures that predict i) mentalizing about the self, ii) mentalizing about another person, and iii) both types of mentalizing. Self-mentalizing and other-mentalizing classifiers had positive weights in anterior/medial and posterior/lateral brain areas, respectively, with accuracy rates of 82% and 77% for out-of-sample prediction. The classifier trained across both types of mentalizing showed 98% predictive accuracy and separated (mental) attributional from factual inferences. Classifier patterns revealed better self/other separation in healthy adults compared to individuals with schizophrenia and with increasing age in adolescence. Together, our findings reveal consistent and separable neural patterns subserving trait-based mentalizing about self and others—present at least from the age of adolescence and functionally altered in severe neuropsychiatric disorders. These mentalizing signatures hold promise as candidate neuromarkers of social-cognitive processes in different contexts and clinical conditions.

**Author Note:** This work was funded by a Starting Grant from the European Research Council (ERC, 101041087) to LKo, a German Academic Exchange Service (DAAD) doctoral grant and a Network of European Neuroscience Schools (NENS) exchange fellowship to DA, an R01 grant from the U.S. National Institutes of Mental Health (R01MH125414-01) to JAH and DAS, a Junior Leader Fellowship from “la Caixa” Foundation (LCF/BQ/PR22/11920017) to PFC, a Consolidator Grant from the European Research Council (ERC, 648082) to LKr, R37, R01 support from the U.S. National Institutes of Mental Health (R37MH076136 to TDW, MH116026 to TDW and L. Chang [PI], R01EB026549 to TDW and M. Lindquist [MPIs]), an NIMH grant (P50MH094258-01A1) to R. Adolphs, and a CIC Brain and Mental Health Chair from the Neurodis foundation to AT. LvdM acknowledges a European Science Foundation EURYI grant (044035001) that funded her doctoral studies (PI: A. Aleman). Views and opinions expressed are however those of the authors only and do not necessarily reflect those of the European Union or the European Research Council. Neither the European Union nor the granting authority can be held responsible for them. The funders had no role in study design, data analysis, manuscript preparation, or publication decisions.

## Introduction

Mentalizing—representing and inferring the psychological states of oneself and others—is a fundamental process for adaptive navigation through the social world^1,2^. Delineating the brain systems involved in mentalizing about self and others is important for understanding brain health and dysfunction, as atypical mentalizing is characteristic for many neurodevelopmental and psychiatric conditions^3–9^. Many studies have examined the neural correlates of mentalizing, suggesting an interplay of different brain regions, including medial prefrontal cortex (mPFC), temporoparietal junction (TPJ), and precuneus^10–14^. However, predictive brain measures of mentalizing that can be easily applied to new data to decode the degree of self-related and other-related processing are still lacking. Most cognitive and affective processes cannot be captured by activity in individual brain regions, as they are reflected in patterns of brain activity distributed across multiple regions and systems, which can be harnessed by pattern-based decoding^15,16^. For instance, recent work demonstrates that distributed brain activity and connectivity patterns, as indexed by fMRI, enable us to decode the intensity of pain^17^, drug and food craving^18^, sustained attention^19^, depressive rumination^20^, and clinically relevant behaviors and outcomes^21–23^. As opposed to the standard brain mapping approach that maps task features onto brain activity, these predictive brain activity patterns or ‘*brain signatures’* are multivariate models that utilize data across the whole brain to make formally testable population-level predictions across subjects and datasets^15,24^. The predictions address the involvement and/or the intensity of a mental process. Here, we apply this brain signature approach to predict mentalizing about oneself or other people.

Mentalizing, as a hidden state, would arguably benefit from such an approach. Thinking about self and others is inherently multilayered and multidimensional^12,25,26^. Recent conceptualizations of self- and social-cognition point to multiple dimensions, such as about oneself versus others, affective versus cognitive^6,12,27^ and multiple layers, such as observing behaviors, and processing at the state and trait levels^25^. Moreover, social cognition terms, including but not limited to mentalizing (e.g., empathy, perspective taking), suffer from inconsistent usage in the literature and a lack of consensual definitions^28^. Developing brain signatures of mentalizing could potentially help account for this heterogeneity, offering the possibility of testing whether the proposed subdimensions of mentalizing are subserved by dissociable neurobiological patterns. In turn, these signatures carry the potential for validating the multiple facets of mentalizing^15^. However, it remains unclear—especially considering the complexity of the mentalizing construct—whether distributed neural patterns can reliably predict mentalizing using brain images.

A central question is whether mentalizing about oneself and mentalizing about others (self- and other-mentalizing hereafter) are reliably distinguishable based on brain activity. Recent electrophysiological evidence suggests that self- and other-mentalizing activate overlapping cortical areas following a similar temporal sequence: TPJ, medial temporal gyrus (MTG)/temporal poles (TP), precuneus/posterior cingulate cortex (PCC), mPFC, in a roughly posterior to anterior temporal order^29,30^. Conversely, differences in brain activity between self- and other-mentalizing have also been observed. In the mPFC, more dorsal areas coincide with other-related processing and ventral areas with self-related processing, with research initially supporting a linear^31^ but more recently a curvilinear^32^ ventral-to-dorsal gradient for self versus other within the mPFC. Further, self-referential thought typically elicits activation in cortical midline structures, such as anterior cingulate cortex (ACC), subcortical areas, including thalamus, striatum and caudate nucleus, and, to a lesser extent, in insula, temporal poles, and ventrolateral prefrontal cortex^31–36^ (vlPFC). In contrast, other-referential thought typically elicits activation in TPJ, middle and superior temporal gyri extending to temporal poles^10,11,32,37–39^, and to a lesser degree in supplementary motor area (SMA), left inferior and medial frontal gyri and medial orbitofrontal cortex^34,40^. Collectively, the literature remains inconsistent regarding the separation between self- and other-mentalizing, and it is unclear whether generalizable brain models of self-versus other-mentalizing can be identified.

To address these gaps, we leverage fMRI and machine learning to develop four distinct brain signatures, 1) the Mentalizing Signature (MS) for mentalizing overall (i.e., thinking about either the self or another person versus non-mentalizing control conditions), 2) the Self-Referential Signature (Self-RS) to detect specifically self-related thought (here referred to as “self-mentalizing”), 3) the Other-Referential Signature (Other-RS) to detect other-related thought (here referred to as “other-mentalizing”), and 4) the Self-vs-Other Referential Signature (SvO-RS) to specifically separate self- and other-related mentalizing. While the overall Mentalizing Signature aimed at developing a broad marker of social reflection that establishes the benchmark ceiling performance and may generalize across a wide range of social processes, the other three signatures (Self-RS, Other-RS, and SvO-RS) allowed us to test whether dissociable and generalizable neural patterns underlie distinct targets of mentalizing.

To this end, we used data from 11 cohorts and seven independent fMRI studies. The training and testing datasets used variants of a standard trait-evaluation task with three conditions, which involved reflecting on personality traits/statements describing oneself or another person. Conceivably, personality traits represent the enduring mental states that can be inferred from information collected by social perceptual systems^41^ across multiple observations^12,25^. Trait representations are used to predict others’ future behavior and become the basis of the “mental-self”^26^. Moreover, trait evaluations are one of the few mentalizing tasks used in the literature to include a self-condition, as other commonly used tasks (e.g., mental attribution, false-belief, perspective-taking tasks) only permit mentalizing about others. Thus, trait evaluation is an ideally suited task to study a key mentalizing process, namely representing stable psychological states of self and others.

We first used standard machine learning algorithms—support vector machines—to develop and cross-validate multivariate classifiers (signatures) of mentalizing about traits in a sample of healthy adults. In a second step, we further validated these signatures in seven independent cohorts from different laboratories, countries, and scanners, and with different sample characteristics (healthy, adolescent, and clinical populations), allowing us to test their generalizability and predictive validity in adolescent and clinical populations. Third, we tested the signatures in two additional independent datasets that used different social cognition tasks to see whether mentalizing signatures would generalize to other social contexts in which mentalizing is likely to occur. Finally, we assessed whether local patterns of brain activity in several regions of interest (ROIs) contain sufficient information to predict self- and other-referential mentalizing. Together, the results of these analyses inform us about the functional neural organization of self-versus other-related mentalizing and provide us with distributed brain signatures that can serve as brain targets for measuring and intervening on mentalizing-related brain processes in future studies.

## Results

### Data overview

The study included a total of 1122 contrast images from 390 participants and 11 independent cohorts, including six samples of healthy adults (*n*=227), two samples of healthy adolescents (*n*=105; *M_age_*=12.9 and 16), and three samples of adults with clinical diagnosis of either schizophrenia (*n*=40) or bipolar disorder (*n*=18; see Fig. 1A and Supplementary Table 1). Thus, this study combined seven independent studies that constituted the training dataset, testing datasets, and the extension datasets (see Fig. 1A). The training and testing datasets included participants completing variants of a self- and other-referential judgement task. The two extension datasets included participants performing a social feedback task and an inference task. Studies 1-5 and 7 included three conditions, namely a Self-condition, an Other-condition, and a nonsocial Control condition, whereas Study 6 included two conditions in one study (attributional versus factual inferences; Study 6a) and a 2×2 design in the second (Study 6b; attributional versus factual inferences x social versus nonsocial scenes). Contrast images were computed for each condition (versus implicit baseline) and rescaled using L2-norm to standardize the scale of beta (regression coefficient) weights across participants, studies, and scanners.

**Figure 1.**
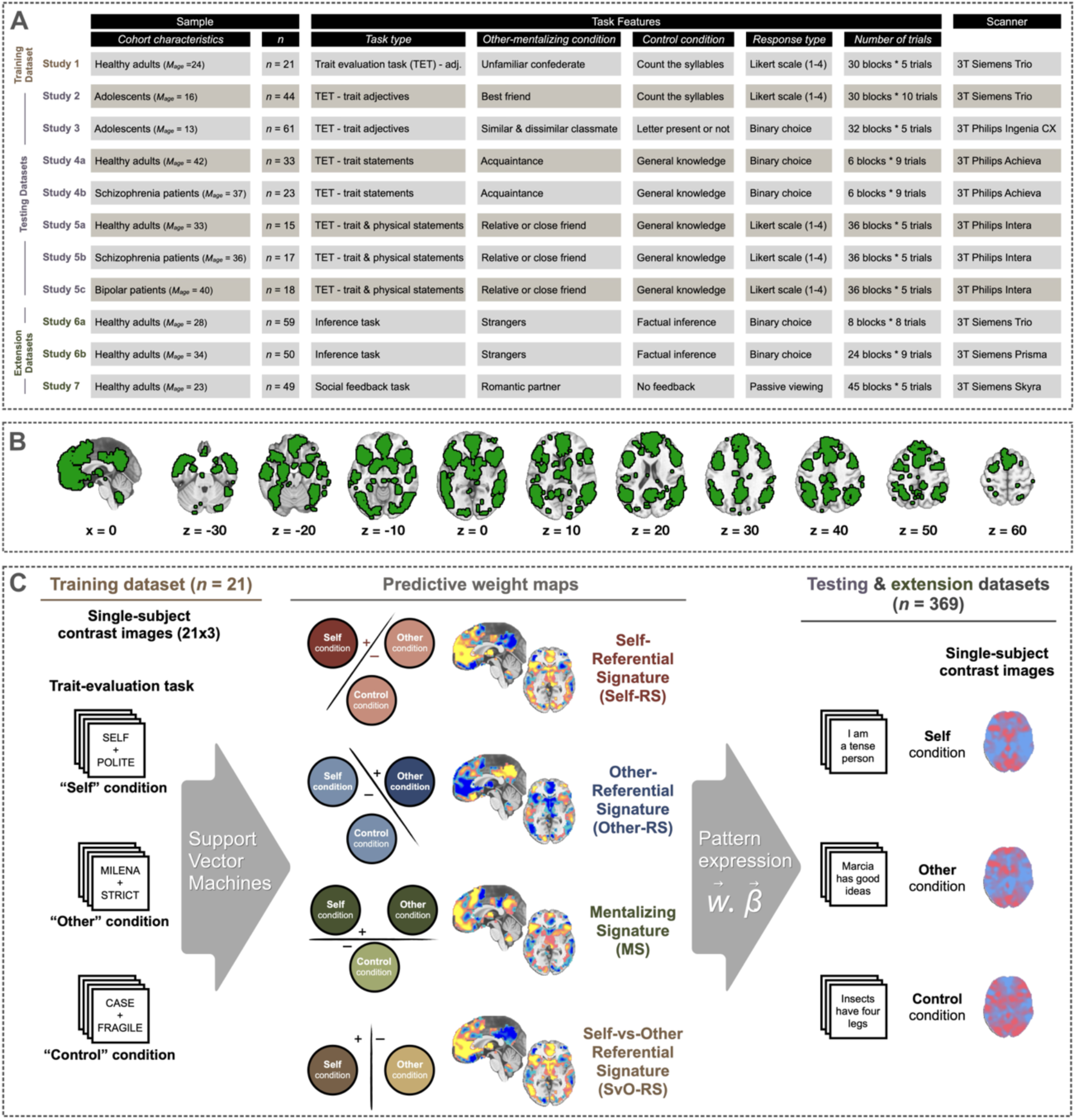
Study design and analytic approach. A) The present study included data from seven independent studies (eleven cohorts, total *N*=390). Training and testing datasets included a trait-evaluation task with three conditions: A Self-condition, an Other-condition, and a nonsocial Control condition. Extension datasets constituted the novel instances to test signatures in distinct tasks. B) Display of the inclusive mask of brain areas related to social cognition. The single-subject contrast images included in this study are masked using this mask. For results with a whole-brain grey matter mask, please refer to Supplementary Fig. 1. C) The training dataset (Study 1) included contrast images of *n* = 21 participants performing a trait-evaluation task with three conditions (Self, Other, Control). We used a 10-fold cross-validated linear SVM to train four mentalizing signatures. The Self-Referential Signature (Self-RS) was trained to predict the Self condition versus the two remaining conditions. The Other-Referential Signature (Other-RS) was trained to predict the Other condition versus the two remaining conditions. The mentalizing signature (MS) was trained to separate the Self and Other conditions from the Control condition. The Self-vs-Other Referential Signature (SvO-RS) was trained to separate the Self and Other conditions. For the validation in independent datasets, we applied the signatures to the single-subject contrast images from Studies 2-7, by computing the dot product between the weight maps and the contrast images.

### Cross-validated training results

The training dataset (Study 1a) consisted of *n*=21 adult participants who completed a trait-evaluation task using a block design (similar to ref^42^; see Fig. 1C). In each block, participants were presented with several trait adjectives: In the Self-reflection condition, they were asked to rate the degree to which each trait adjective described themselves. In the Other-reflection condition, they had to rate how much each adjective described another person—a confederate with whom the participant had previously interacted during a decision-making task^43^. In the non-mentalizing Control condition, participants indicated the number of syllables in the trait adjectives. To reduce the influence of non-mentalizing-related brain regions (e.g., visual cortex) and opportunistic classification based on features not related to mentalizing, we used an inclusive mask of brain regions related to social processing (see Fig. 1B).

We used support vector machines (SVM) with default parameters (to avoid overfitting) and 10-fold cross-validation^44^ to train three distinct mentalizing signatures using the masked contrast images from the training dataset. The folds were defined at the subject level, so that all contrast images from a given subject were assigned to the same fold. The Self-Referential Signature (Self-RS) was trained to detect Self-reflection (versus the two remaining conditions), the Other-Referential Signature (Other-RS) to detect Other-reflection (versus the two remaining conditions), the Mentalizing Signature (MS) to detect both types of mentalizing versus the Control condition, and the Self-vs-Other Referential Signature (SvO-RS) to differentiate Self-versus Other-related processing (see Fig. 1C).

All four signatures showed excellent cross-validated (out-of-sample) prediction accuracy (100% accuracy in two-alternative forced-choice tests, *p*<.001, *Cohen’s d* for the Self-RS = 3.18, the Other-RS = 2.45, the MS = 4.92; the SvO-RS = 2.45; AUC = 1.00 for all signatures). To identify which voxels contributed most reliably to the mentalizing signatures, we used bootstrapping (5000 samples) to obtain one *p*-value per voxel and displayed the thresholded weight maps using false discovery rate (FDR) correction at *q* < .05 and cluster extent *k* > 10 voxels. Brain regions with significant positive voxel weights for the Self-RS (see Fig. 2) included the ventromedial (vmPFC) and dorsomedial (dmPFC) prefrontal cortex, frontal eye fields, ventral ACC, frontal operculum, anterior insula, thalamus, and caudate nucleus (see Supplementary Table 2). For the Other-RS, significant positive weights were found in left vlPFC, left superior temporal sulcus (STS), bilateral TPJ, and precuneus/PCC (see Fig. 2 and Supplementary Table 3). For the SvO-RS (see Fig. 3 and Supplementary Table 5), brain regions with significant positive voxel weights (thus, associated more with the Self than the Other condition) included vmPFC/ACC, left anterior insula, left inferior frontal sulcus, left caudate nucleus, left middle temporal cortex, and bilateral thalamus. Significant negative clusters in the SvO-RS (thus, associated more with the Other than the Self condition) were found in precuneus, bilateral TPJ/inferior parietal cortex, right PCC, left STS, and left inferior frontal gyrus. The MS had significant positive clusters in mPFC (both dorsal and medial), bilateral vlPFC, dorsal ACC (dACC), frontal operculum, bilateral SMA, bilateral MTG, bilateral TP, left STS, middle cingulate gyrus (MCC), bilateral PCC, precuneus, bilateral angular gyrus/TPJ, and subcortical areas, including right caudate nucleus, left anterior insula, and bilateral thalamus (see Fig. 2 and Supplementary Table 4).

**Figure 2.**
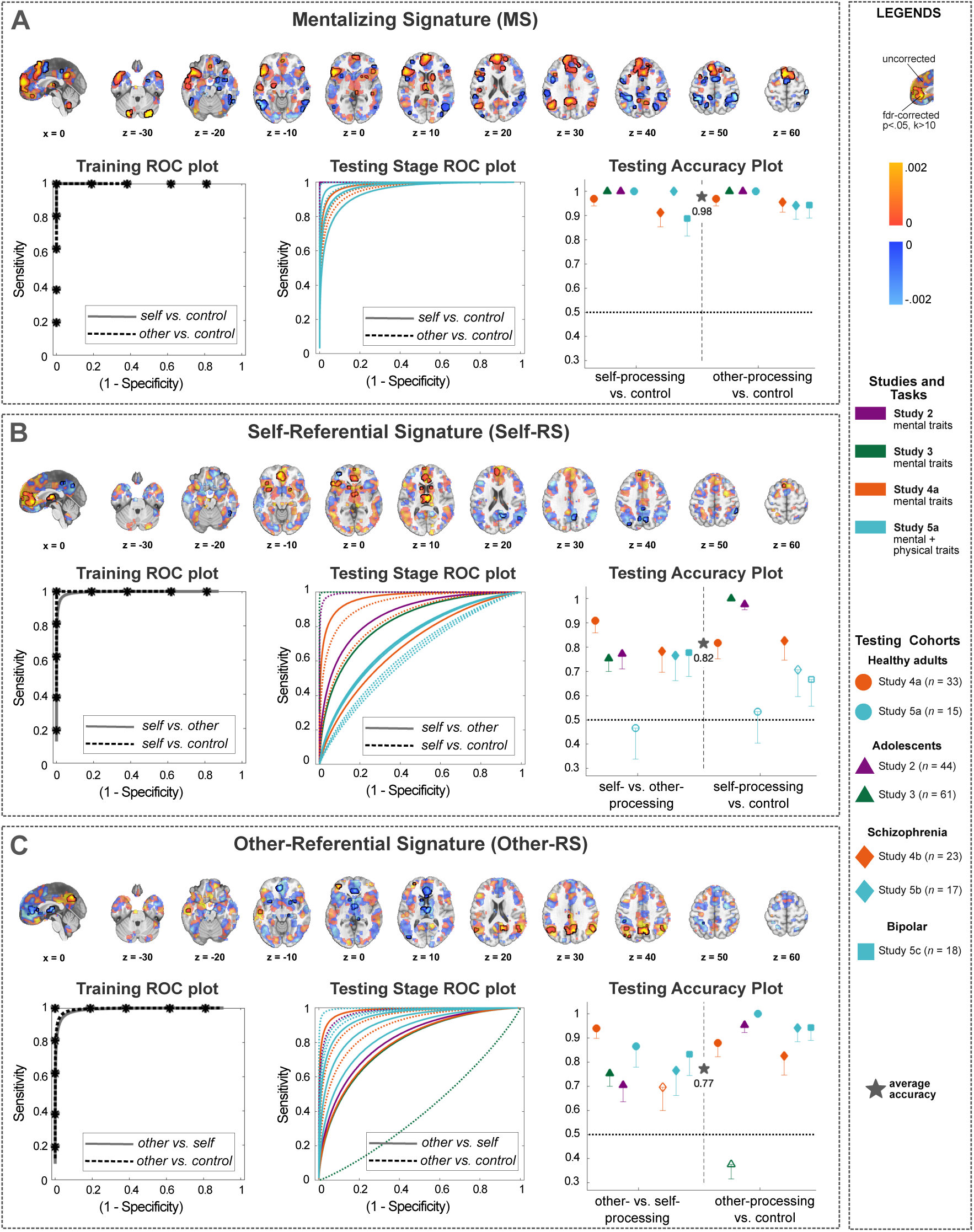
Model weights, cross-validated training, and validation results of the mentalizing signatures. On the top row of each box, weight maps illustrate the positive and negative weights of A) the Mentalizing Signature (MS), B) the Self-Referential Signature (Self-RS), and C) the Other-Referential Signature (Other-RS). Voxels significant at an FDR-corrected threshold (*q* < .05, minimum cluster size of *k* = 10) are highlighted by black outlines and brighter colors. The left and middle plots in each box depict the receiver operating characteristic (ROC) plots from the i) cross-validated training and ii) testing datasets. The last column shows the accuracies of the mentalizing signatures in each testing dataset separately, with color representing the study, and shape illustrating the cohort type. The one-sided error bars below shapes illustrate the standard error. The star shows the weighted average accuracy across all seven datasets. To ensure that our cross-validated training and validation results were not influenced by the social-processing mask that we used, we repeated our analyses with the whole-brain (grey-matter masked) images, (see Supplementary Fig. 1). To test if a different feature selection method would yield similar results, we trained classifiers using recursive feature elimination (RFE), which we report in Supplementary Fig. 2.

**Figure 3.**
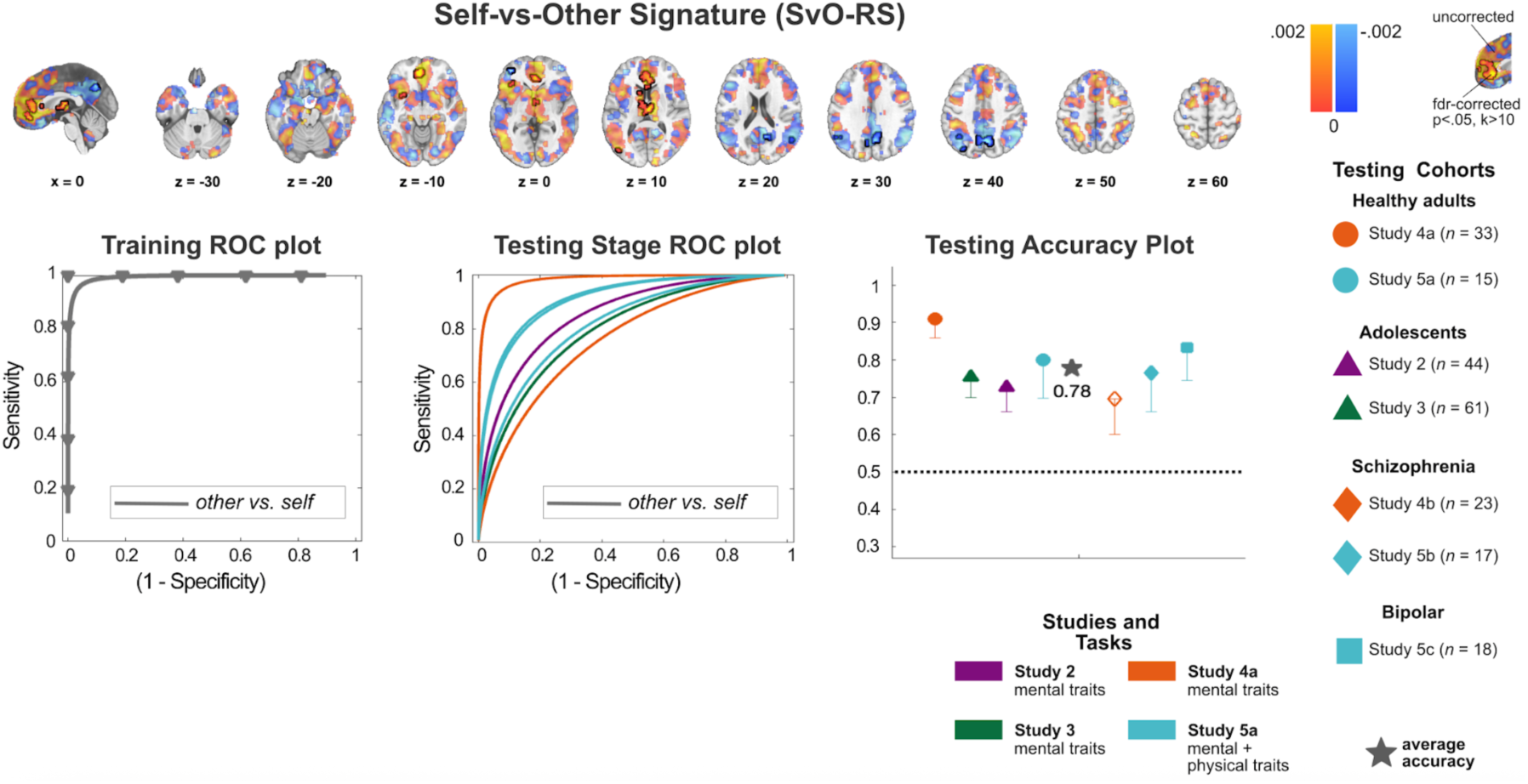
Model weights, cross-validated training, and validation results of the Self-vs-Other Classifier (SvO-RS). The weight map (first row) illustrates the positive and negative weights, as well as the voxels significant at an FDR-corrected threshold (*q* < .05, minimum cluster size of *k* = 10). The second row depicts the receiver operating characteristic (ROC) plots from the i) cross-validated training and ii) validation stages, and iii) the accuracies of the SvO-RS in each testing dataset. The cohort types are shape-coded, and the studies are color-coded in the validation accuracy plot. The one-sided error bars below shapes illustrate the standard error. The star shows the weighted average accuracy across all seven datasets.

### Validation in independent samples

Next, we validated the four brain signatures on several completely independent studies: two samples of healthy adults (Studies 4a & 5a), two adolescent samples (Studies 2 & 3), one cohort of participants with bipolar disorder (Study 5c), and two cohorts of participants with schizophrenia (Studies 4b & 5b). All datasets included comparable trait evaluation tasks and fMRI block designs with three conditions, with some small variations in task designs between studies (see Fig. 1A). To obtain pattern expression values, we computed the matrix dot product between the mentalizing signatures and each subject-level contrast image from these datasets, yielding one scalar value per individual contrast image and per signature (see Fig. 1C). These pattern expression values were then used to test the predictions of the mentalizing signatures. The weighted average prediction accuracy in two-alternative forced-choice (2AFC) tests, across all independent testing datasets, was 81.52% for the Self-RS (+/- 6.17% average STE; significant in 10 out of 14 validation tests [7 samples*2 comparisons]), 77.25% for the Other-RS (+/- 6.02% average STE; significant in 12 out of 14 tests), 97.87% for the MS (+/- 1.79% average STE; significant in all 14 tests), and 77.73% for the SvO-RS (+/- 7.14% average STE; significant in 6 out of 7 tests), suggesting overall high prediction accuracy of the four signatures, even in new samples, although with some variations between datasets (see Fig. 2, Fig. 3 & Supplementary Table 6). There were no sex effects on pattern expression or signature fit—defined as the difference between the pattern expression for the true condition and the false condition in each signature (see Methods).

### Lower self/other separation in participants with schizophrenia

The results above show that the signatures significantly predicted mentalizing in most samples, including clinical samples. Yet, schizophrenia, in particular, is often associated with impaired social cognition and altered self-perception^45,46^. Thus, we next tested how well the signatures separated self-from other-related mentalizing in two of the testing datasets (Study 4 & Study 5) that included both participants with schizophrenia (total *n* = 40) and matched healthy participants (total *n* = 48). As expected, the three signatures, the Self-RS (*M_HC_* = .32, *M_SCZ_* = .12; β = .21, *STE* = .07, *CI* = [.07, .35], *t*(86) = 2.99, *p* = .004), the Other-RS (*M_HC_* =.43, *M_SCZ_* = .24; β = .19, *STE* = .07, *CI* = [.04, .33], *t*(86) = 2.57, *p* = .012), and the SvO-RS (*M_HC_* =.39, *M_SCZ_* = .19; β = .20, *STE* = .07, *CI* = [.07, .34], *t*(86) = 2.97, *p* = .004) showed better discrimination between self- and other-referential mentalizing in healthy adults, compared to adults with schizophrenia (i.e., a greater positive difference between the Self-RS responses in the Self compared to the Other condition, and a greater difference between the Other-RS responses for the Other compared to the Self condition). No group differences were found in the correct discrimination of Mentalizing versus Control by the MS (*p* = .29, see Fig. 4). These results indicate that, compared to healthy adults, participants with schizophrenia have less differentiated brain patterns between self- and other-related thought, indicating the potential clinical utility of the signatures.

**Figure 4.**
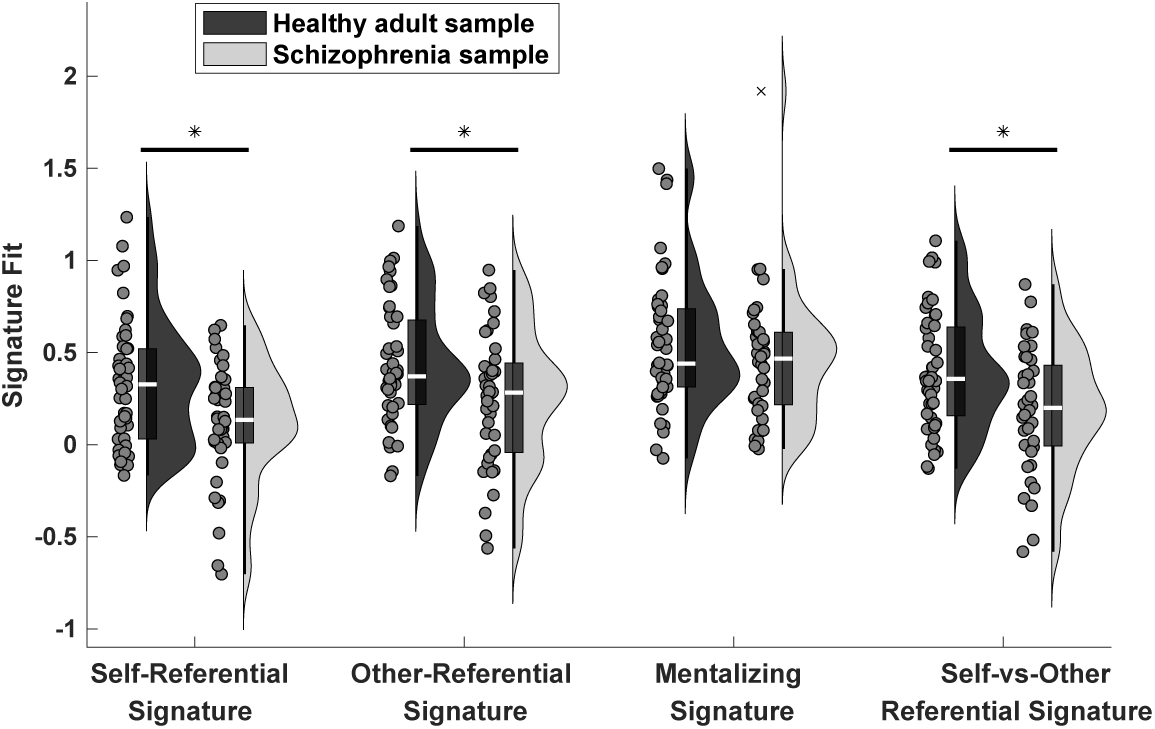
Self/other discrimination based on classifier responses in healthy adults compared to individuals with schizophrenia. Signature fits are measured as the pattern expression of true minus false class, separately for each signature in healthy adults (*n*=48, in dark grey) and adults with schizophrenia (*n*=40, in light grey). Self/other discrimination was found to be better in Self-RS, Other-RS, and SvO-RS. Linear mixed effects are used to test the group differences, controlling for different cohorts. Self-RS: *β* = .21, *STE* = .07, *CI* = [.07, .35], *p* = .004. Other-RS: β = .19, *STE* = .07, *CI* = [.04, .33], *p* = .01, SvO-RS: β = .20, *STE* = .07, *CI* = [.07, .34], *p* = .004. Dots indicate values per person, and boxplots mark the median, lower, and upper quartiles. Outliers (> 3 SD from the mean) are marked with an “×”. Whiskers extend from the minimum to the maximum data points, excluding outliers. Asterisks indicate statistical significance: *p < .05.

For completeness, we also compared the discrimination of self-versus other-related mentalizing activity in participants with bipolar disorder (*n* = 18) versus healthy adults (*n* = 15) using data from Study 5. However, participants with bipolar disorder did not differ significantly from controls for any of the signatures.

### Better self-other separation with increasing age in adolescents

Next, we explored potential developmental differences in the classifier performances by testing whether the ability of the classifiers to separate self-from other-related mentalizing depended on the age of the participants in the two adolescent samples (*N* = 105). We combined data from Study 2 (age 12-18 years; *M_age_* = 16) and Study 3 (age 11-14 years; *M_age_* = 12.9). We found that (controlling for sex and the source study) older adolescents had better self/other separation for the three signatures trained to distinguish these two conditions: the Self-RS (β = .08, *STE* = .03, *CI* = [.018, .14], *t*(102) = 2.55, *p* = .012), the Other-RS (β = .06, *STE* = .03, *CI* = [.001, .12], *t*(102) = 2.03, *p* = .045), and the SvO-RS (β = .07, *STE* = .03, *CI* = [.01, .13], *t*(102) = 2.31, *p* = .023; see Fig. 5). In contrast, there were no significant associations between age and performance of the MS. These results suggest that, with increasing age, adolescents’ brain activity becomes more differentiated for self-versus other-related mentalizing.

**Figure 5.**
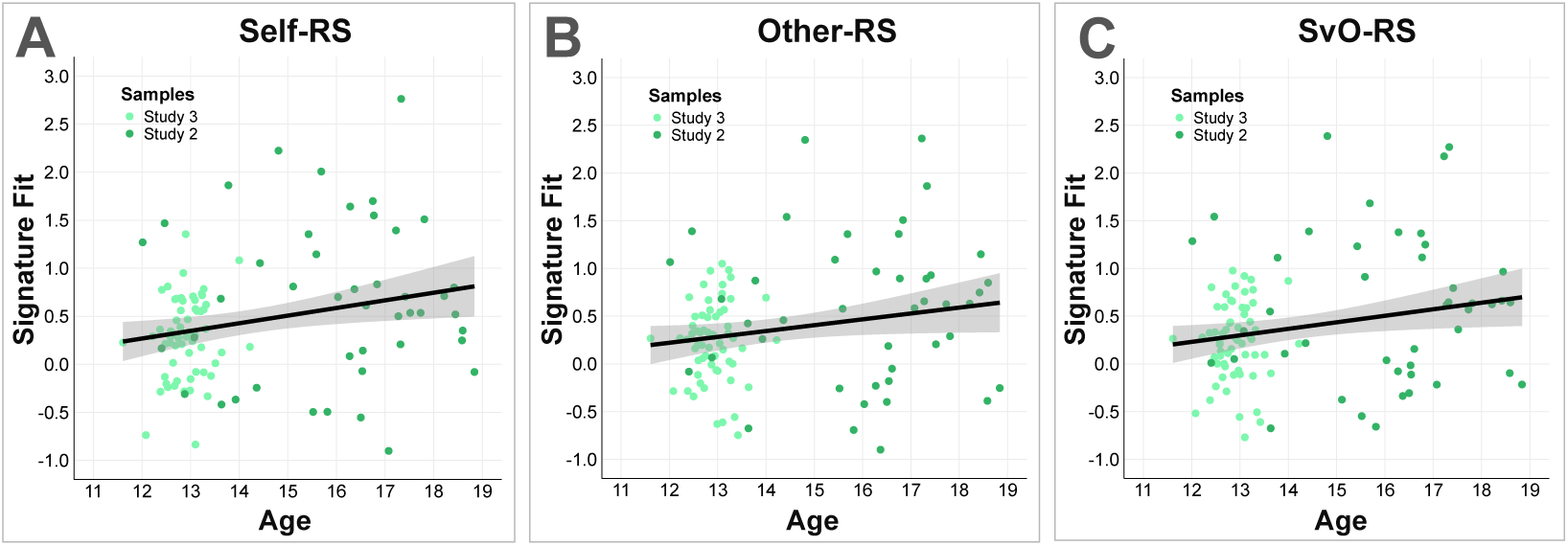
Association between self/other discrimination and age of adolescents (n = 105). Older age correlated significantly with better differentiation of self- and other-related mentalizing, controlling for cohort (Self-RS [β = .08, STE = .03, CI = [.012, .018], p = .01]; Other-RS [β = .06, STE = .03, CI = [.001, .12], p = .047]; SvO-RS [β = .07, *STE* = .03, *CI* = .009 to .12, *p* = .02]. Black lines display the linear regression fits, shaded areas the 95% confidence intervals. Dots show values per participant.

### Testing the signatures during attributional versus factual inference

So far, we have shown that the signatures performed well in several independent datasets from different labs, but all using a similar explicit mentalizing task. In order to test whether signatures are implicated specifically in mentalizing and not in broader social or even nonsocial reflections, we tested them in an additional dataset (Study 6a & 6b) of n=109 healthy adults making attributional versus factual inferences about social and nonsocial stimuli (see Fig. 6A for the task). Attributional inferences (e.g., “Is Ada proud of herself?”), especially in a social context, require mentalizing, or in other words, they require representing hidden mental states. Factual inferences (e.g., “Is Ada tall?”), on the other hand, refer to directly observable features and should not require mentalizing. Therefore, we tested whether the Other-RS and the MS could predict attributional against factual inference conditions successfully.

**Figure 6.**
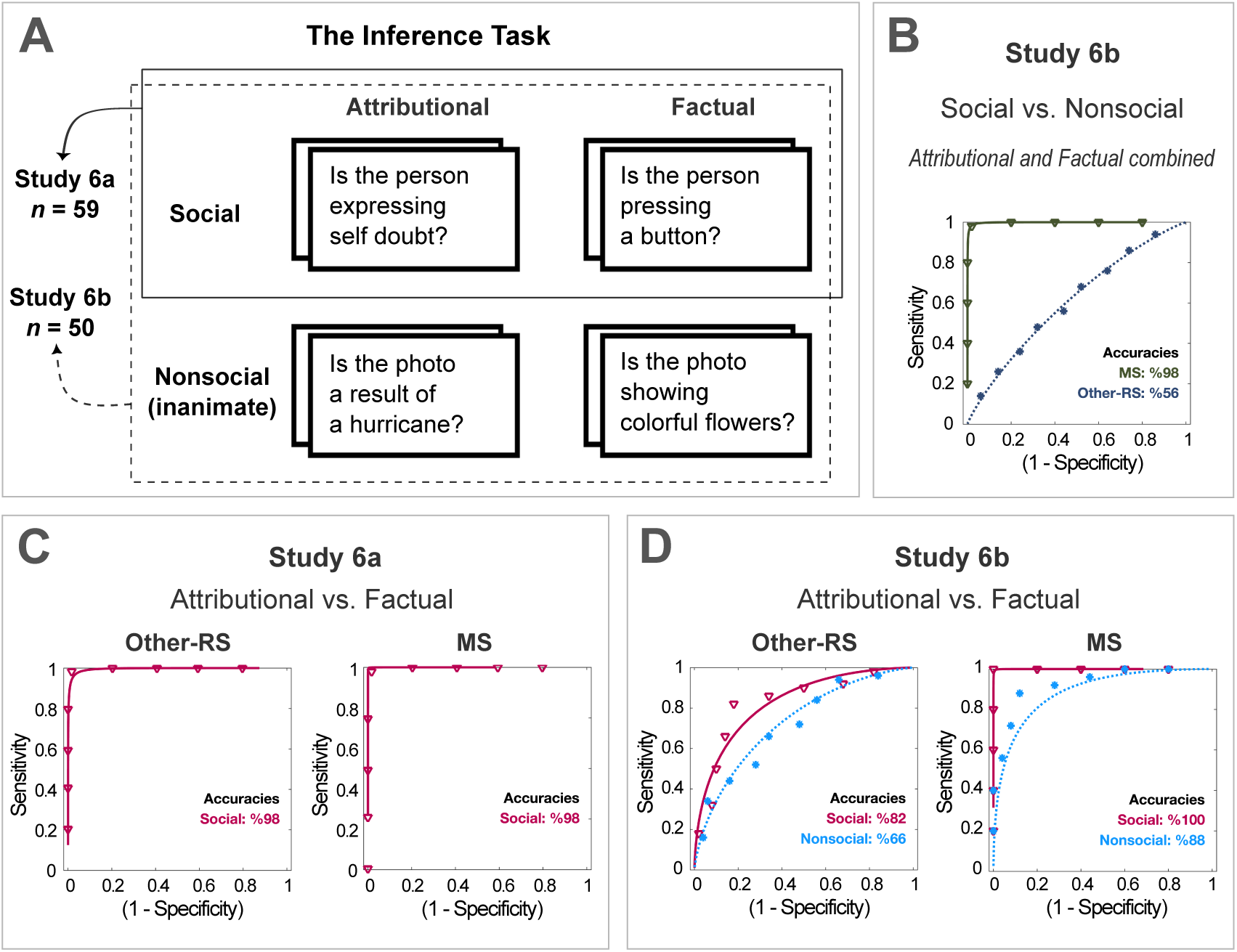
Extension of the Other-RS and the MS to an independent inference task (total n=109). A) Task design. Study 6a had a 1×2 design (attributional versus factual inferences regarding social scenes). Study 2 had a 2×2 design (attributional versus factual inferences x social versus nonsocial scenes). B) Binary classification performances of the MS and the Other-RS in social versus nonsocial attributions (Study 6b). C) Binary classification results from Study 6a for separating attributional versus factual inferences. D) Binary classification results from Study 6b, suggesting that the Other-RS and MS signatures are more sensitive to attributional (versus factual) inferences that require reflection about internal states—especially those concerning social agents.

Participants in Study 6a (*n* = 59) were asked to make these two types of inferences on social images showing faces or hands with a question prompt (e.g., is the person proud of themselves?). Attributional (versus factual) inference conditions were predicted with 98% accuracy (2AFC) by both the Other-RS and the MS (see Fig. 6C; *p* < .001, AUC = 1.00, sensitivity = 98%; specificity = 98%, *Cohen’s d* = 2.62 for the Other-RS and 3.94 for the MS). Study 6b had a 2 (attributional versus factual) x 2 (social versus nonsocial) design. The Other-RS predicted the attributional inferences against factual inferences with 82% accuracy in the social condition (*p* < .001, *Cohen’s d* = .95, AUC = .85, sensitivity = 82%, specificity = 82%) and 66% accuracy in the nonsocial condition (*p* = .03, *Cohen’s d* = .61, AUC = 0.73, sensitivity = 66%, specificity = 55%; see Fig. 6D). The MS predicted the attributional versus factual inference with 100% accuracy in the social condition (*p* < .001, *Cohen’s d* = 3.56, AUC = 1.00, sensitivity = 100%, specificity = 100%) and 88% accuracy in the nonsocial condition (*p* < .001, *Cohen’s d* = 1.21, AUC = 0.92, sensitivity = 88%, specificity = 88%; see Fig. 6D). Further, we tested how well the signatures separate social from nonsocial inferences (averaged across attributional and factual conditions). While the Other-RS did not significantly separate social from non-social inferences (56% accuracy, *p* = .48, Cohen’s *d* = 0.27, AUC = 0.62, sensitivity = 56%, specificity = 56%), the MS separated social from nonsocial inferences with almost perfect accuracy (98% correct, *p* < .001, Cohen’s *d* = 2.96, AUC = 1.00, sensitivity = 98%, specificity = 98%; see Fig. 6B). Together these results suggest that the mentalizing signatures are sensitive to processes that require a reflection about internal or hidden states, and–at least for the MS–especially for social agents.

### Testing the signatures in a social feedback task

To examine how the signatures would respond to social interaction tasks in which mentalizing is not directly instructed but likely implicated, we next tested their performance in an fMRI dataset (Study 7) of *n*=49 romantic partners performing a social feedback task. Participants were led to believe that the task was designed to understand how people rate other people and interpret others’ ratings. Therefore, participants first indicated how likable they felt each of 20 individuals (confederates) was, who supposedly rated the participants and their partners in return. To set up a context facilitating spontaneous mentalizing, participants were told that their partners saw the ratings on the screen simultaneously, in a separate room. The task included conditions in which participants received feedback about their own likeability (self-feedback), the likeability of their partner (partner-feedback), as well as a no-feedback control condition. The control condition consisted of trials where no feedback was given, with a prior explanation that such trials signaled that the rater (the individual on the screen) was not shown the participants’ picture or did not supply their rating in the time required by the rating task.

Here, we tested whether the signatures’ responses paralleled the target of the feedback conditions. For instance, to provide evidence of successful extension, the Self-RS should show higher pattern expression values in the self-feedback condition as opposed to the other two conditions. As expected, the Self-RS responded more strongly to the Self-condition, *F*(2,96) = 14.214, *p* < .001, η_p_^2^ = .23 (Fig. 7). Bonferroni-corrected pairwise comparisons showed that the Self-feedback condition (*M* = .16) had significantly higher Self-RS expression scores than both the Partner-feedback (*M* = −.09, *p*=.01) and Control (*M* = −.36, *p* < .001) conditions. Additionally, the Self-RS also produced higher pattern expression values for the Partner-feedback (*M* = −.09) compared to Control condition (*M* = −.36, *p* = .04). Similarly, the MS successfully discriminated both Feedback conditions against Control condition, *F*(2,96) = 14.711, *p* < .001, η_p_^2^ = .24 (Fig. 7). Bonferroni-corrected pairwise comparisons indicated that both Self-feedback (*M* = .17, *p* < .001) and Partner-feedback (*M* = .12, *p* = .002) conditions had significantly higher pattern expression scores than the Control condition (*M* = −.17). However, the Other-RS did not produce any significant differences between task conditions, *F*(2,96) = 2.003, *p* = .14 (Fig. 7). Together this suggests a modulation of the MS and the Self-RS (but not the Other-RS) even in a task where mentalizing was not directly instructed, in line with the idea that feedback to oneself or one’s partner should lead to more mentalizing related brain activity.

**Figure 7.**
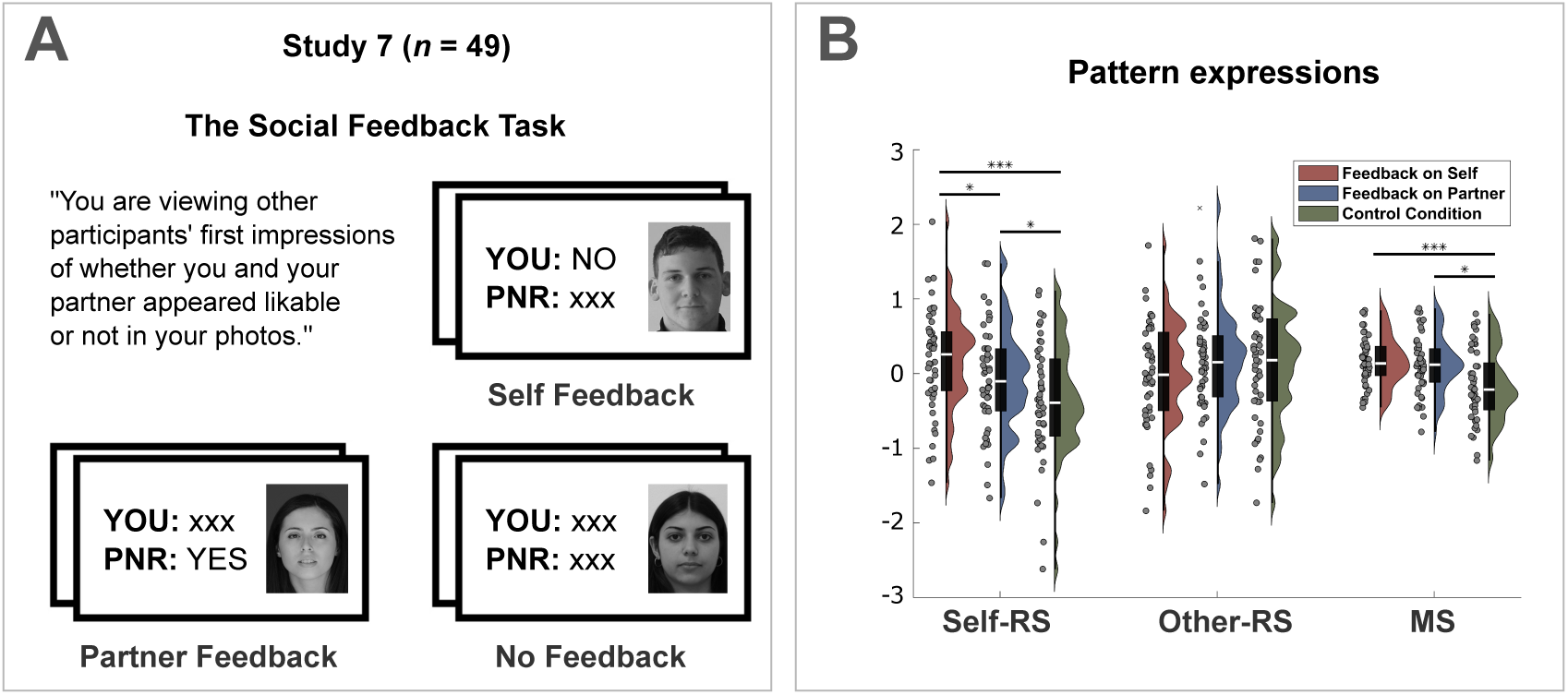
Extension of three signatures to an independent social feedback task (n=49). A) Participants received feedback for themselves, for their partners, or no feedback (Control condition). B) Signature responses for each of the three conditions, demonstrating higher Self-RS responses to feedback concerning oneself than the partner and compared no feedback, and higher mentalizing-related brain responses (MS) for self- and partner-directed compared to no feedback. Dots indicate values per person, and boxplots mark the median, lower, and upper quartiles. Outliers (> 3 SD from the mean) are marked with an “×”. Whiskers extend from the minimum to the maximum data points, excluding outliers. Asterisks indicate significant Bonferroni-corrected pairwise comparisons following an initial repeated measures ANOVA test within each signature: *p < .05; ***p < .001.

### Local patterns of self- and other-related mentalizing

We trained local classifiers in regions of interest (ROI) associated with mentalizing to gain further insight into how mentalizing-related information is processed locally and which regions can predict self-versus other-related mentalizing. The whole brain patterns indicate which regions have the most reliable positive and negative contributions to different forms of mentalizing, but they do not necessarily show which regions are *not* involved and do not inform us whether local patterns alone can predict the target of mentalizing (self or other). Training and testing classifiers for brain regions implicated in the literature can inform us in this regard.

To this end, we included ten ROIs (see Fig. 8) previously associated with mentalizing as shown in an automated term-based meta-analysis (NeuroSynth^47^): mPFC, two clusters in anterior middle temporal gyrus (aMTG) bilaterally, two clusters in TPJ bilaterally, a cluster covering precuneus and posterior cingulate cortex (precuneus/PCC), left SMA, and three clusters in the cerebellum. We trained four types of classifiers in each of these ten ROIs. In other words, we trained each ROI to perform the four following classification tasks separately: 1) Self-mentalizing (versus the two remaining conditions); 2) Other-mentalizing (versus the two remaining conditions), 3) both mentalizing conditions (self and other) against the Control condition, and 4) Self-mentalizing against Other-mentalizing. The ROI classifiers were trained and cross-validated in our training dataset (Study 1a; *n* = 21) and tested in the combined sample (*n* = 211) of participants from all testing datasets (including all healthy, developmental, and clinical cohorts; for detailed results, see Supplementary Table 7).

**Figure 8.**
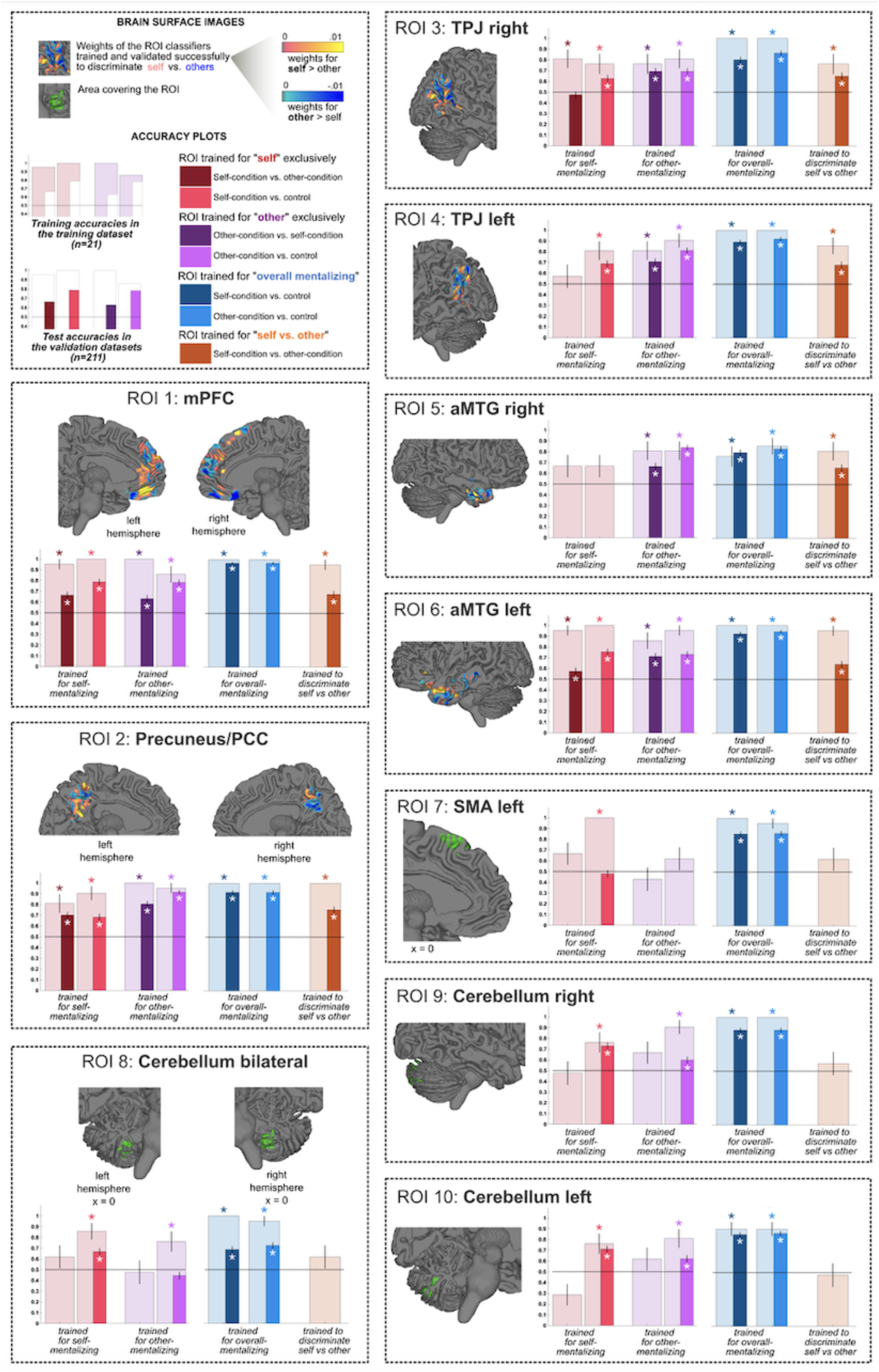
Results of the ROI analysis. Cross-validated training and validation results for ten ROIs derived from a term-based meta-analysis (‘mentalizing’, NeuroSynth^47^): mPFC, precuneus/posterior cingulate cortex (precuneus/PCC), temporoparietal junction (TPJ) right, TPJ left, anterior middle temporal gyrus (aMTG) right, aMTG left, supplementary motor area (SMA) left, cerebellum right, cerebellum left, and a bilateral cerebellum cluster. The ROIs that were consistently predictive of both self- and other-related mentalizing were: mPFC, precuneus/PCC, TPJ, aMTG. Four different classifiers were trained for each ROI: (i) to predict Self exclusively, (ii) to predict Other exclusively, (iii) to predict overall mentalizing, and (iv) to discriminate the Self versus Other. If an ROI classifier had significant accuracy in the cross-validated training dataset, it was further tested across the testing datasets. The cross-validated training accuracies are illustrated by the wider bars in the background. The validation accuracies are illustrated by the thinner bars at the front. The error bars represent the standard error of the prediction. Classification performances significantly above the chance level (50%) are marked with asterisks. The surface images illustrate the weights of the classifiers that were trained to discriminate Self versus Other. Orange-yellow colors show areas with positive weights towards Self, blueish colors show areas associated with positive weights towards Other. If a specific ROI did not yield significant training and independent classification accuracy for this task, then the area covering this ROI is displayed in green color.

As expected, all ten regions showed excellent classification accuracy as the general mentalizing classifiers in the training dataset, and all predicted mentalizing successfully in the testing dataset (see blue bars in Fig. 8). However, not all regions could significantly differentiate Self-reflection and Other-reflection conditions. While mPFC, aMTG, TPJ, and precuneus/PCC were capable of differentiating self and other conditions, the cerebellum clusters and left SMA failed at this task (see orange bars in Fig. 8). When trained for self-mentalizing (see red bars in Fig. 8), the right aMTG cluster could not capture any consistent configurations for Self-mentalizing. While TPJ subregions were successful here against control conditions, they could not predict Self-against Other-mentalizing, implying a lack of exclusive neural patterns for self-mentalizing within TPJ subregions. Finally, when trained for Other-mentalizing, the Cerebellum subregions and left SMA were incapable of picking up consistent patterns (see violet bars in Fig. 8).

Together, our analyses suggest that mPFC (including both vmPFC and dmPFC), TPJ, aMTG, and precuneus/PCC clusters encode both self- and other-mentalizing, and that the left SMA and Cerebellum clusters most likely are involved in domain-general operations during mentalizing. Besides, our findings suggest that TPJ and aMTG are more strongly associated with other-mentalizing, while mPFC and precunes/PCC clusters are unique in the sense that they are consistently involved in both self- and other-related mentalizing.

### Leave-One-Site-Out classifiers trained on all datasets

Finally, to ensure that the high accuracies were not an artifact of features unique to the training study (e.g., scanner, sample), we trained new classifiers that combined data across all five training and testing datasets using a leave-one-site-out (LOSiO) cross-validation approach. We combined images from *n* = 232 participants from Studies 1 to 5 that used a trait-evaluation task, pooling data from healthy participants, adolescents, and clinical samples. We used the same parameters to train SVM models corresponding to the four mentalizing signatures.

Cross-validated prediction accuracies of all four classifiers were very good to excellent (see Fig. 9A-D): 83% averaged accuracy (2AFA) for the Self-RS_LOSiO_, 91% for the Other-RS_LOSiO_, 96% for MS_LOSiO_, and 90% for the SvO-RS_LOSiO_ in two-alternative forced-choice tests, *p* < .001 (averaged *Cohen’s d* for the Self-RS_LOSiO_ = 1.10, for the Other-RS_LOSiO_ = 1.36, for the MS_LOSiO_ = 0.64, for the SvO-RS_LOSiO_ = 1.19; averaged AUC for the Self-RS_LOSiO_ = 0.90, for the Other-RS_LOSiO_ = 0.95, for the MS_LOSiO_ = 0.93, for the SvO-RS_LOSiO_ = 0.93). Thus, these results show that the high prediction accuracies achieved by the mentalizing signatures were not inflated by task-specific idiosyncrasies but rather reflect consistent, robust multivariate patterns that are achieved using different combinations of the training dataset.

**Figure 9.**
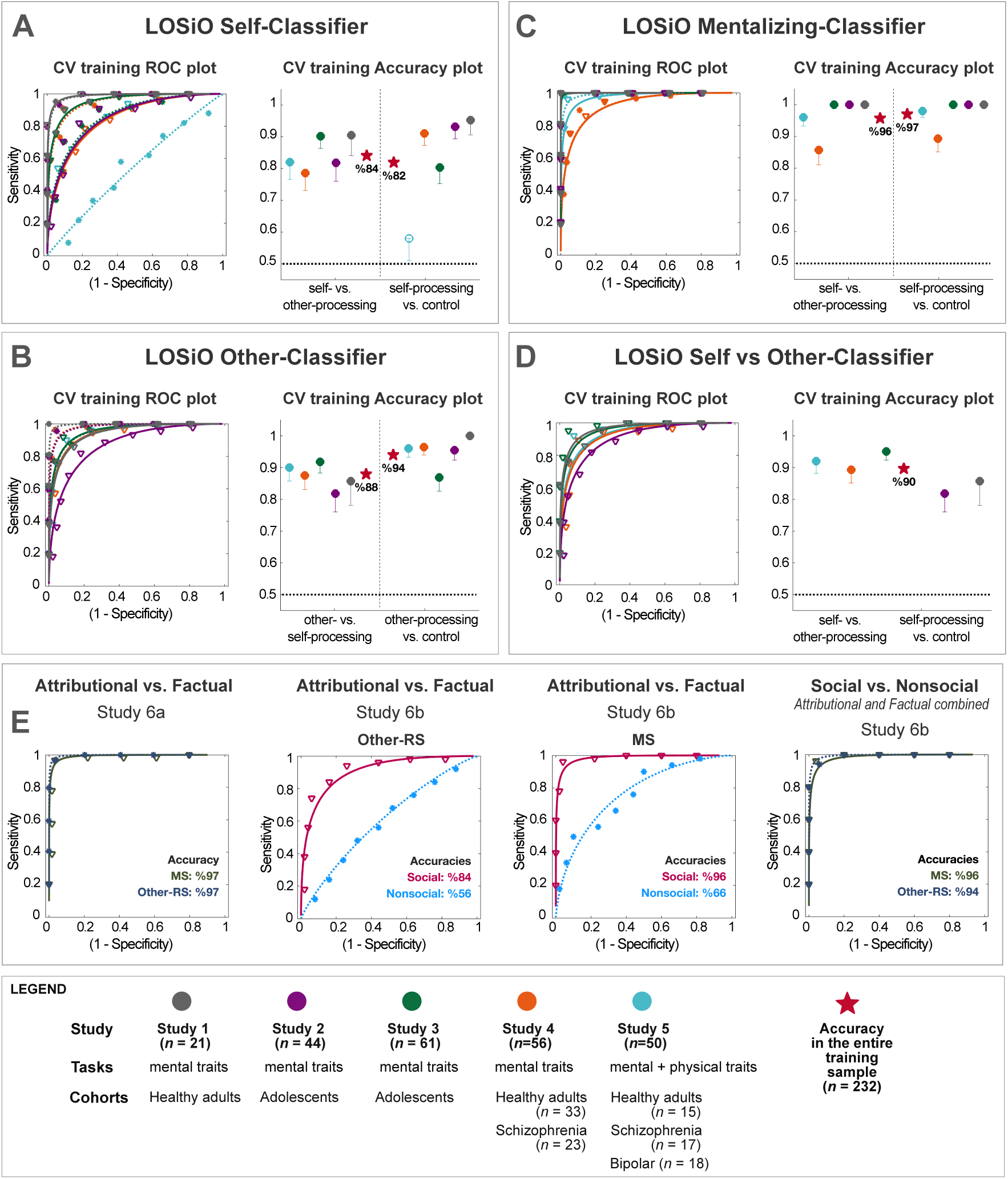
Leave-one-site-out (LOSiO) classifiers. We trained additional classifiers that combined data across all datasets that used a trait-evaluation task (n=232) to rule out that the results were driven by idiosyncratic features of the original training dataset. The classifiers were trained using the same analytic strategy as the main mentalizing signatures, but using five folds, with each study being held out in one-fold. A) Self-classifier, B) Other-classifier, C) Mentalizing-classifier, D) Self-vs-Other classifier. Red stars show the out-of-sample accuracy across the whole sample (n=232), while the colored circles represent the cross-validated accuracy for each study when classifiers were trained on all other studies’ data. The one-sided error bars in accuracy plots illustrate the standard error. E) ROC plots for classifications in the inference task (Study 6) that involve attributional versus factual statements in social and nonsocial settings.

Finally, we applied the predictions of the LOSiO classifiers to the inference task (Study 6) to test how well they classify attributional versus factual and social versus nonsocial conditions (see Fig. 9E). Both Other-RS_LOSiO_ and MS_LOSiO_ predicted social attributional (versus factual) statements in Study 6A with 97% accuracy (*p* < .0001; Cohen’s *d* for Other-RS_LOSiO_ = 2.75 and for MS_LOSiO_ = 2.42). In Study 6b, the Other-RS_LOSiO_ separated attributional versus factual statements in the social condition with 84% accuracy (*p* < .001, Cohen’s *d* = 1.32), whereas this classification was non-significant in the nonsocial condition (accuracy = 56%, *p* = .48). The MS_LOSiO_ predicted attributional versus factual statements in the social condition with 96% accuracy *p* < .001, Cohen’s *d* = 2.16) and in the nonsocial condition with 66% accuracy (*p* = .033, Cohen’s *d* = 0.73). Finally, MS_LOSiO_ and Other-RS_LOSiO_ separated social from non-social inferences with 96% and 94% accuracy respectively (*p* < .001, Cohen’s *d* for MS_LOSiO_ = 2.12, and for Other-RS_LOSiO_ = 2.5). Together, this illustrates that, similar to the mentalizing signatures, the LOSiO classifiers are most sensitive to attributional processing in the social context, while also capturing nonsocial attributional processing to some degree.

## Discussion

Mentalizing—reflecting about others’ and one’s own internal states—is a fundamental capacity required to function in a social world. Transdiagnostically, mentalizing deficits typify an array of psychopathological conditions^3–9^. Here, we developed novel brain signatures of self- and other-related mentalizing (*Self-RS*, *Other-RS, and SvO-RS*), as well as mentalizing in general (the ‘*Mentalizing Signature’* or *MS*), combining data from eleven independent cohorts (*N*=390). The four signatures showed good to excellent classification accuracies in independent data and provided several insights into the brain circuits of trait-based mentalizing. First, mentalizing about traits appears to coincide with a specific pattern of brain activity, with reliably dissociable activation patterns for self- and other-related mentalizing, based on both whole-brain activity as well as within key nodes of the social brain, especially mPFC, precuneus/PCC, TPJ and aMTG. Second, the signatures significantly predicted mentalizing not only in healthy adults, but also in adolescents and in individuals with schizophrenia and with bipolar disorder, demonstrating its utility across different (including clinical) samples. Third, our results point to the relevance of these mentalizing signatures for a range of different research questions in social, developmental, and clinical neuroscience, by showing (i) engagement of the signatures in independent social cognition tasks, (ii) that the differentiation of self- and other-related mentalizing increases in transition from adolescence to adulthood, and (iii) that it is less pronounced in individuals with schizophrenia compared to healthy controls. Thus, our data provide us with clearly defined *brain models*^15^, applicable across contexts to assess the engagement of mentalizing-related brain regions in different experimental conditions and their alteration in clinical and neurodevelopmental conditions.

The Mentalizing Signature (MS) successfully predicted mentalizing across different contexts and extended to tasks without explicit mentalizing instructions (e.g., spontaneous mentalizing). It performed very well in separating higher-level inferences about others and hidden states from factual inferences, with higher classification in social compared to non-social contexts. The MS (but not the Other-RS) also significantly separated social from nonsocial inferences. Together, these findings demonstrate its sensitivity to mental state compared to external state attributions. The regions that contributed to the MS most positively included mPFC, precuneus, TPJ, STS, and many other regions previously associated with mentalizing^10,13,29,38,48^, providing convincing face validity of this signature. Its high predictive performance in testing datasets implies that mentalizing recruits consistent and reliable neural configurations in the brain, not only in healthy controls, but also in clinical cohorts and developing adolescents. The MS also extended to other tasks (social-feedback and attributional versus factual inferences), predicting conditions that potentially evoke implicit, spontaneous mentalizing. This suggests that implicit and explicit forms of mentalizing largely share a common neural basis, in keeping with previous research^49^. Thus, subnetworks of mentalizing (implicit-explicit) may be building on a common anatomical architecture that is here covered by the MS.

Recent work proposes that self- and other-mentalizing activate common neural constellations or representations and recruit largely overlapping regions^29,30,48^. In contrast, our findings suggest that, while they share a large common core (as evident in the MS), they can also be distinguished reliably based on both whole-brain and local activation patterns. The Self-RS and the SvO-RS had positive weights in anterior mentalizing regions around cortical midline structures, such as mPFC, ACC, thalamus, caudate nucleus, frontal eye fields, and insula, extending to the frontal operculum. This pattern is in line with previous meta-analyses^31,34,36^. The mPFC has been consistently related to mentalizing, and especially its ventral part, with self-related processing^33,39,42,50,51^. The ventral-to-dorsal linear^31^ and curvilinear gradient^32^ hypotheses propose that mPFC is involved in both self-and other-person related processing. Accordingly, our results show that mPFC is involved in both self- and other-related mentalizing with different neural configurations, and that the information encoded in mPFC seems to predict self-mentalizing more consistently. Besides, self-processing recruited reward-processing circuits (e.g., striatum), in line with the idea that introspection about oneself is intrinsically rewarding^52–54^. The involvement of subcortical areas (e.g., thalamus) and cortical areas associated with interoception (e.g., insula) is in line with previous work suggesting that one has access to affective/physiological information during self-mentalizing, which are less pertinent during mentalizing about others^55^, and that interoceptive information might be an important contribution to one’s sense of self^26,56,57^.

The Other-RS and (negative weights of) the SvO-RS subsumed more posterior and lateral regions, such as TPJ, precuneus/PCC, STS, and vlPFC, resonating with previous findings^10,11,13,32,37–39^. Interestingly, our results concerning the precuneus/PCC do not align with the known subdivisions of the default-mode^58^ or mentalizing networks^30^. This region is thought to be part of the midline core of the default-mode network^58^ and of the medial subsystem of the mentalizing network^30^, with stronger associations with self-processing. However, here in our results, while the precuneus/PCC cluster also contains information about self-mentalizing, it appears more strongly associated with other-related mentalizing, along with the areas that are part of the lateral subdivisions of both networks (e.g., TPJ, temporal lobes). A potential explanation here could be the involvement of precuneus in mental orientation/imagery^59,60^.

The mentalizing signatures generalized to data from different types of cohorts, including healthy adults, schizophrenia and bipolar samples, and adolescents. This shows that the overall neurobiological configurations of self- and other-related mentalizing are comparable across these populations and that the signatures have utility in different types of study populations. However, the Self-RS, the Other-RS, and the SvO-RS signatures also showed sensitivity to clinical status, since self/other differentiation in the pattern responses was significantly less pronounced for both patterns in participants with schizophrenia compared to matched healthy controls. This finding is noteworthy and in line with clinical observations of alterations in self-perception, mentalizing, and the ability to discriminate between self- and other-generated thought^36,45,46,61^. The responses in the bipolar sample did not differ significantly from those of the healthy adults but given the limited sample size of the bipolar group in the present analysis, future work is needed to assess more fine-grained (and potentially context-dependent) differences in mentalizing responses in bipolar disorder, as well as in other psychiatric and neurodevelopmental disorders with mentalizing deficits.

In the two adolescent samples (aged 11 to 18 years), greater age was associated with better self/other differentiation of the Self-RS, the Other-RS, and the SvO-RS. In other words, responses of these brain signatures to self- and other-related mentalizing were more different from each other in older adolescents. Considering that the signatures were developed using an adult dataset, this reflects an ongoing development of the mentalizing neurobiology^62,63^ to become more adult-like and more differentiated for different targets of mentalization, during the age span covered in our study (11-18 years). Future studies could test the role of pubertal development instead of chronological age, and test the signatures in younger children, and those with developmental disorders.

Of note, the mentalizing signatures were also implicated in making inferences about nonsocial targets—albeit to a lesser degree compared to social targets (see Fig. 6D and Fig 9E). An earlier study^64^ that used a similar inference task as Study 6 found that social and nonsocial attributions displayed overlapping activations in dmPFC, STS, TPJ, and lateral orbitofrontal cortex. A common denominator in both conditions may reflect domain-general attributional processing^65^, which at least partially builds on semantic cognition^66^. Likewise, processes supporting mentalizing may also more broadly subserve extraction of conceptual and semantic meaning for nonsocial stimuli^67,68^, in line with the default mode network’s putative meaning-making function^51,69,70^. Thus, the mentalizing signatures may involve components of both domain-general representational processing, domain-specific social processing, and target-specific (self and other) processing components.

We note that the signatures’ classification performance in the cross-validated training is consistently higher than in the independent testing datasets. This could potentially be due to leakage of information across the different participants in the training dataset, who all performed the same task in the same scanner and experimental settings. However, the additionally developed leave-one-site-out (LOSiO) classifiers that combined data across five studies performed similarly to the brain signatures that are developed using data from only the training dataset (Study 1). This provided evidence that the results of this study are not an artifact of our analytic choices and that the methodological approach fits the aims of the study well, as the results are robust with different permutations. The LOSiO classifiers showed consistently high classifications for the original training dataset (Study 1), which further provided support for the use of this study for training. Yet, because the LOSiO classifiers used information from n=232 participants during model development, it has potential to generalize very well to independent contexts. A key feature of the brain signature approach^15^ employed here is that the brain models are not necessarily final products, but tunable, improvable working models. Therefore, we endorse the use of LOSiO classifiers in future research along with the main mentalizing signatures for further evaluation of their utility in diverse settings.

The high separability of self- and other-referential processing in Study 1 might also reflect the fact that the Other-condition in the mentalizing task was about a relatively unfamiliar person (a confederate). This may lead to a greater difference in self-versus other-related processing compared to studies in which mentalizing was about a familiar or close person—which is often more closely related to the self and likely also engages self-referential processing. This could potentially also explain why the Other-classifier was less successful in separating other-related from self-related feedback in the sample of romantic couples (Study 7), since participants may consider their partners as an extension of themselves. Relatedly, the lower performance of signatures in Study 5 (compared to other testing datasets) could be due to the use of physical attributes (in addition to personality traits), which rely less on mentalizing.

Of note, the mentalizing signatures developed here primarily capture trait-related mentalizing. Here, we provide initial evidence that they extend beyond explicit reflections on trait-level representations. However, they also show above-chance classification in nonsocial inferences (see Fig 6D and Fig. 9E), which might raise concerns about their specificity to social agents. Future studies are needed to further test how well the mentalizing signatures separate mentalizing about other agents from other higher-order, conceptual, and abstract cognitive processing. Future studies should also test their specificity, as well as their validity in other forms (e.g., affective, spontaneous, or non-verbal) of mentalizing. A natural next step could be to test whether other dimensions of mentalizing^6,12^ (e.g., cognitive versus affective) or different types of mental state content^71^ (i.e., stable versus transient or beliefs versus preferences) are also associated with robust, replicable multivariate patterns. Recent work^72,73^ investigated valence and self-relevance as two key components of internal thought. It is an open question how the brain integrates valence with different targets during mentalizing. Finally, future research can also build on this work by testing the signatures in other mentalizing tasks (e.g., false-belief, emotion imagery), on a greater variety of targets^74^, and in other related mental processes (e.g., autobiographical memory retrieval).

In conclusion, we trained and validated four brain signatures that predict self-related, other-related, and both types of mentalizing in multiple independent datasets that used different variants of a standard trait-mentalizing task in diverse populations. Our findings imply that self- and other-mentalizing use largely dissociable neural mechanisms that build on a foundational overall mentalizing capacity. Indeed, the four mentalizing signatures possess potential for use as predictive neuromarkers of mentalizing about self and others, for example by testing how brain-circuits of mentalizing are engaged or modulated by different types of experimental conditions, and how they might be altered in different psychiatric and neurodevelopmental populations.

## Methods

### Participants

The present study pooled data from a total of *N*=390 participants (119 females and 162 males, *M_age_*= 25.5, *SD_age_*= 12.9) from eleven cohorts participating in seven independent studies (see Fig. 1A and Supplementary Table 1). These eleven cohorts comprised six samples of healthy (neurotypical) adults, two samples of healthy adolescents, and three samples of adults with a diagnosed psychiatric condition. While most of the datasets have been previously published separately, the analyses reported here were not published previously, and the seven studies have not been previously combined.

The training sample (Study 1^43^) consisted of *n*=21 healthy adults (10 women and 11 men, *M_age_* = 23.5) who were recruited at the University of Geneva, Switzerland. One additional participant with structural abnormalities in the brain was excluded from the original study and the present analysis.

Study 2^75^ included *n*=44 healthy adolescents (23 females and 21 males, *M_age_* = 16, *SD_age_* = 1.86, age range = 12.01 to 18.84) who were recruited from secondary schools in Geneva, Switzerland. In the original study, one additional participant was excluded due to structural abnormalities in the brain, three due to non-completion of the paradigm, one due to signs of substance use, and five due to excessive movement.

Study 3^76^ included *n*=61 healthy adolescents (27 females and 34 males*, M_age_* = 12.9, *SD_age_* = 0.43, age range = 11.61 to 14.22) who were recruited for a longitudinal project from secondary schools in the Netherlands. An additional 18 participants were excluded from the original study due to excessive movement, incorrect task completion, or measurement errors. Additionally, 5 participants were not included in this study (original study n = 66) because they did not provide explicit consent for data sharing.

Study 4^77,78^ included n=33 healthy adults (14 women and 19 men, *M_age_* = 41.7) and *n*=23 adults with schizophrenia (7 women and 16 men, *M_age_* = 37). The two cohorts were matched on age, sex, and a measure of general intelligence. The patients were recruited from a local psychiatric hospital in Barcelona, Spain, based on diagnostic interviews. The schizophrenia diagnosis was confirmed using the Structured Clinical Interview for DSM Disorders^79^ (SCID).

Study 5^80^ included three types of cohorts: Healthy adults (*n*=15, 6 women and 9 men, *M_age_* = 33.3), adults with schizophrenia (SZ, *n*=17, 6 women and 11 men, *M_age_* = 35.5), and adults with bipolar disorder (BD, n=18, 9 women and 9 men, *M_age_* = 40.3). The clinical cohorts were recruited from a local hospital in the north of the Netherlands. The diagnoses of the patients were confirmed using the Mini International Neuropsychiatric Interview-Plus^81^ 5.0.0 (MINI-Plus). All BD patients were chosen among those who had a history of at least one psychotic episode. All three cohorts were matched with one another on age, sex, level of education, and a measure of general intelligence. The SZ and BP patients were additionally matched on the level of cognitive and clinical insight as measured by the Schedule of Assessment of Insight^82^-Expanded version (SAI-E, clinical insight) and the Beck Cognitive Insight Scale^83^ (BCIS, cognitive insight).

Study 6a^84^ included 59 healthy adults (26 female and 33 male; *M_age_*= 28.3), and Study 6b^84^ included 50 (19 female and 31 male; *M_age_*= 33.6) healthy adults from the Los Angeles metropolitan area; these final sample sizes reflect exclusions described below. One participant from each study was excluded due to poor task performance (>70% missed trials), and four additional participants from Study 6b were excluded due to outlier behavioral scores (>3 SDs from the mean, n=1) and excessive scanner motion (n=3). All participants were right-handed, had normal or corrected-to-normal vision, spoke English fluently, and had IQs in the normal range (assessed via the WASI-II).

Study 7^85^ included 56 healthy adults in romantic relationships recruited from the Tucson, Arizona community and surrounding areas. Seven participants were excluded from the study due to missing or inadequate imaging data, yielding a final sample size of *n*=49 (26 women and 23 men, *M_age_* = 22.6). Community members were eligible to participate if they had been in a romantic relationship for at least six months, had no contraindications for MRI scanning, and did not meet criteria for active psychosis or mania at the time of screening. Both members of the couple completed all components of the study, including the social feedback fMRI task.

All participants gave written informed consent and were compensated for their participation via monetary means or gifts. All studies were approved by the respective institutional ethics committees. For additional information, please refer to the original studies and Supplementary Table 1.

### Tasks

#### Training dataset

We used a trait evaluation task, adapted from ref^42^ as the training mentalizing task. Participants were asked to rate the extent to which certain personality adjectives (e.g., ‘talkative’, ‘daring’) described themselves (Self-condition) or a same-sex confederate (Other-condition), with whom they thought they were interacting in a preceding decision-making task^43^. In the Control condition, they were asked to count the syllables for each trait adjective.

Participants met the confederate in person at the beginning of the experimental session, where they were briefed that one of them would be tested in the scanner, the other one in a separate room, and that they would interact via computer interface. They first participated in a social decision-making task resembling a serial dictator paradigm, whereby the participant was given the choice to share or keep the resources with the confederate in a series of trials^43^. After this task, participants were introduced to the trait-evaluation task and were asked to rate their co-player.

The task consisted of 150 trials in total, presented in 30 blocks (ten per condition). Each block contained five trials and lasted for 20 seconds, with an inter-block interval (fixation cross) of 8 seconds. Half of the blocks only contained negative and the other half only positive adjectives. Each word appeared once for each condition (50 adjectives were used in total).

Each trial started with a 3-second cue above the fixation cross, reminding the participant of the condition (self, other, or syllables), which was followed by the presentation of the adjective and a Likert scale (1-4) on which the participant was expected to select using a button press device. A word was presented every 4 seconds. If the participant made a choice sooner, the word would disappear for the remaining time of the 4 seconds before the next trial. The order of the blocks and words displayed in each block was randomized.

#### Testing and extension datasets

All testing datasets included similar mentalizing tasks with the same three conditions (Self, Other, Control) as the training task and utilized a block design (for an overview, see Fig. 1A and Supplementary Table 1). The main difference between validation tasks was the stimuli that were presented: Studies 2 and 3 used trait adjectives, Study 4 (with two cohorts) used trait statements, and Study 5 (with three cohorts) used a mix of trait and physical statements. The studies also used a variety of *others* in the Other-condition: Best/close friend (Study 2), similar and dissimilar classmate (Study 3), relative or close friend (Study 5), and acquaintance (Study 4). Studies 2 and 5 collected responses using Likert scales, and Studies 3 and 4 asked for binary choices (yes/no). Finally, the Control conditions also varied between studies. Studies 4 and 5 presented general knowledge statements; Study 3 asked participants to search for the letter in words; Study 2 asked them to count the syllables in words.

Extension datasets (Studies 6 and 7) used different tasks to test how well the mentalizing signatures extend to tasks that involve other forms of social (or nonsocial) processing. Studies 6a and 6b used an inference task based on visual stimuli^84^ (see Fig. 6A). In Study 6a, participants made rapid yes/no judgments in response to social scenes (intentional hand actions and emotional faces) that required either attributional inferences about hidden/internal states (mentalizing) or factual inferences about observable features, resulting in a 1 (stimulus type: social) × 2 (inference type: attributional, factual) factorial design. Study 6b included an additional nonsocial condition using images of natural scenes (e.g., rain pouring out of a gutter), each paired with attributional or factual inference prompts, resulting in a 2 (stimulus type: social, nonsocial) × 2 (inference type: attributional, factual) design. Study 7 (see Fig. 7A) investigated spontaneous empathy when negative or positive feedback thought to be from other participants (confederates) was directed to participants’ romantic partners (or one’s self). Participants received positive and negative feedback either about themselves (self-condition), their romantic partners (other-condition), or no feedback (control-condition). The original study (see ref^85^ for more details) included more conditions, such as directing feedback to both self and partner, which are not analyzed in the present manuscript.

### fMRI data acquisition, preprocessing, and first-level analysis

#### Training dataset

The training MRI images were acquired on a 3T Magnetom TIM Trio whole-body scanner (Siemens, Germany) with the product 12-channel head coil. A T1-weighted MPRAGE sequence (TR = 1900ms, TI = 900 ms, TE = 2.27 ms, voxel size 1 x 1 x 1 mm) was used to acquire structural anatomical images. Functional images were obtained using a standard T2-weighted echo-planar imaging sequence (2D-EP, TR = 2100 ms, TE = 30 ms, flip angle 80°, voxel size 3.2 x 3.2 x 3.2 mm) that scanned the whole brain in 36 sequential slices. An automated shimming procedure was included to minimize magnetic field inhomogeneities.

As it is the standard practice in the development of brain signatures^15,17,18,21^, we used statistical maps from first-level analysis (e.g., single-subject contrast images) as input for training and testing the classifiers. SPM8 (Wellcome Department of Imaging Neuroscience, UCL, London, UK) and Matlab® (The MathWorks Inc.) were used for image preprocessing and first-level analysis. A standard preprocessing pipeline was performed, which included spatial realignment and reslicing of the functional images, coregistration, unified segmentation and normalization to the Montreal Neurological Institute (MNI) echo planar imaging template (voxel size: 2 mm^3^), and finally spatial smoothing using an 8 mm Full Width at Half Maximum (FWHM) Gaussian kernel.

During first-level analysis, we included six task (block) regressors that were composed of 3 task conditions by positive and negative valence. The task regressors were convolved with a canonical hemodynamic response function. We also included six additional regressors of no interest to control for motion. A high-pass frequency filter (128s) and autocorrelation corrections (using restricted maximum likelihood and an autoregressive model) were used in model estimation.

#### Testing datasets

We conducted a non-systematic literature review to identify recent fMRI studies of comparable mentalizing tasks with at least three conditions (Self, Other, and non-mentalizing Control condition) and in which participants rated trait adjectives or statements. We emailed the authors of seven studies that we identified and received positive responses from four of them. Data from these four studies were included in the current study as testing datasets. In addition, to test the signatures in a different type of task, we included an unpublished dataset^85^ (Study 7) and data from recently published work^84^ (Study 6) as extension datasets. Please refer to the original studies (see Supplementary Table 1) for details of image acquisition, preprocessing, and first-level analysis.

The authors provided the subject-level contrast images of the different experimental conditions (versus the implicit baseline). Voxel weights were normalized within each image using L2-norm, and contrast images were resampled onto the same image space as the training dataset using linear resampling. For Study 3, which included two different Other-conditions (similar and dissimilar classmates), we averaged the contrast images across these two conditions, resulting in a single Other-condition, as in the training and the other testing datasets (analyzing them separately did not alter the results). Study 6 included two types of social conditions: hands and faces. As we were not interested in the social processing of hand- or face-related information, we averaged the contrast images of these conditions, creating a single Social Condition image per participant.

## Data Analysis

### Cross-validated training

Using 10-fold cross-validation, we trained three support-vector-machine (SVM) classifiers in Study 1 that discriminate each condition from the other two conditions: The Self-RS was trained to separate the Self-condition from Other and Control conditions, the Other-RS to separate the Other-condition from Self and Control conditions, and the MS was trained to separate both mentalizing conditions (Self and Other) from the Control condition. In addition, we trained one signature to specifically separate the Self from the Other condition (the SvO Signature). The input for SVMs was subject-level contrast images for each of the three task conditions (versus implicit baseline). The cross-validation folds were organized to ensure that the three contrast images from each participant were kept together within a single fold (nine folds with two participants and one fold with three participants). To reduce the possibility that classifiers opportunistically used non-mentalizing related processes (e.g., visual information), we applied a mask in the training dataset that includes key social-cognition regions (see Fig. 1B). This mask was computed as the union of six term-based meta-analytic maps (association and uniformity maps for “mentalizing”, “self-referential”, and “social”, downloaded from NeuroSynth^47^ on 06/06/2024) to cover a wide range of brain areas typically activated by social and self-related processing. While the use of a mask was intended to minimize involvement of the confounding voxels that are not directly related to social cognition, this might raise the question of how more extensive, whole-brain classifiers might perform. As shown in Supplementary Fig. 1, applying a whole-brain gray matter mask yielded highly similar results.

SVMs were chosen based on their high performance for binary linear classification problems in high-dimensional and low-sample data^86–88^. Linear SVM is a commonly used method in cognitive neuroscience in decoding problems^48,89,90^, which is computationally efficient and provides interpretable, robust weight maps. The algorithm fits a hyperplane that classifies true and false classes by assigning weights for each feature (i.e., each voxel). The fitting is performed for each of 10 folds, in which the data of 90% of participants are used for fitting the classifier, and the resulting classifier is tested on the remaining 10% hold-out participants’ data, allowing for assessing its classification performance in independent hold-out data. Because using a one versus the rest approach in SVM (e.g., Self versus Other and Control, see Fig. 1C) may add bias into the model by favoring the majority class, we fitted weighted SVM models with a ridge parameter of .5 to train the Self-RS, the Other-RS, and the MS (but not the SvO-RS). To avoid overfitting, the SVMs were otherwise trained using default parameters (regularization parameter *C* = 1, linear kernel function, number of folds = 10). The cross-validated distance from the hyperplane of hold-out images was used to calculate the receiver operating characteristic (ROC) curves and the accuracy for each classification. Each SVM results in a weight map with one value per voxel. These voxel weights are effectively the predictive weights of the true class, yielding brain signatures of mentalizing that can be applied to other brain images to obtain a single pattern expression score per image. The classification accuracies in the training dataset were tested using two-alternative forced-choice predictions and binomial tests.

We used bootstrapping to train SVM classifiers, producing stable model estimates and voxel-wise *p*-values. In the first step, cross-validated training generated prediction accuracies and a final model output (i.e., weight map) that is trained on all samples. In the next step, the bootstrapping procedure resampled the training data 5000 times with replacement, yielding statistical maps with two-tailed, uncorrected p-values. These maps were then thresholded using FDR-correction at *q* < .05 with a minimum cluster size of 10 voxels. Note that corrected maps were only used for visualization and inference. The unthresholded weight maps constitute the mentalizing signatures and, therefore, are used for analysis.

### Validation in independent datasets

The four mentalizing signatures were applied to the testing datasets by computing the pattern similarity values as the matrix dot product between mentalizing signatures and the subject-level (1st level) contrast images of each study, yielding one scalar value per condition and participant and signature. The predictions followed a binary forced-choice principle using paired observations. The signatures’ classification accuracies were assessed using Receiver Operating Characteristic (ROC) analysis and binomial tests using a two-sided significance threshold of *p* < .05. The accuracy rate, area under the curve (AUC), and specificity/sensitivity obtained from these analyses provide the benchmark measures of binary classifications in multivariate modeling^44^. We also report Cohen’s d in the text as a more standardized measure to quantify prediction effect sizes across studies and samples. Cohen’s d is calculated as the mean of differences between condition values (either pattern expression or distance from the hyperplane) divided by the pooled standard deviation of differences (where difference = input_values[binary_class] - input_values[∼binary_class]). Because the three signatures (Self-RS, Other-RS, and MS) were each tested in two comparisons (e.g., Self versus Other and Self versus Control), the main text reports the average statistical values across these comparisons, with the complete results provided in the Supplementary Tables. Similarly, when accuracy statistics from a classifier were combined across conditions or studies, we report the weighted average values in the main text and present the full set of statistics in the Supplementary Material.

To examine the effects of individual differences (e.g., sex and age) on pattern expression values, we used linear mixed-effects models. We pooled data from five testing datasets that used the same task (*n* = 211; Studies 2–5). The signature fit (“true minus false class score”) was defined as the difference between the pattern expression for the true condition and that for the false condition. Specifically, we subtracted the false-condition value from the true-condition value for each signature: Self-RS (Self – Other), Other-RS (Other – Self), SvO-RS (Self – Other), and MS ([Self + Other]/2 – Control). Age and sex were entered as fixed effects, and study was included as a random intercept in the mode: *fit ∼ age + sex + (1 | study)*.

#### Group comparisons of pattern expression values

To quantify how well signatures discriminate true versus false conditions and to easily compare self/other separation across groups, we used the signature fits for each signature, the calculation of which is explained in the section above. For statistical comparisons between participants with schizophrenia and healthy controls, we ran linear mixed effects models for each of the signatures, with the signature fits (‘true minus false class score’) as the dependent variable. Group constituted the fixed effect variable (healthy controls [*n*=48] versus schizophrenia sample [*n*=40]). Because the data came from two different studies (Study 4 and Study 5), study was included as a random effect variable. Therefore, the model equation was “*signature_fit ∼ group + (1|study)*”.

To test whether the bipolar sample (*n*=18) differed from healthy adults (*n*=15) in Study 5 regarding their pattern expression values, we performed independent samples t-tests for each of the signatures consecutively. The dependent variable was the “signature fit” as above.

#### Associations with age

We tested whether the adolescents’ age was associated with self-other discrimination using linear mixed effects models. We included age as the fixed effect, Study (Study 2 and Study 3) as the random effect, and sex as the control variable (fixed effect). The dependent variable was the difference of the true minus false classes for each of the signatures, as detailed above. Therefore, the model formula was “*signature_fit ∼ age + sex + (1|study)*”.

### Testing the signatures in extension tasks

In order to test how well the signatures generalized to the two extension tasks, we applied them to the subject-level contrast images for each experimental condition by computing their dot products. When a signature is applied to another context (e.g., a different task), one could apply either i) a binary prediction test or ii) a linear-effects model to test for condition differences, depending on the study aim. Binary prediction is arguably most appropriate when there is a ground truth or clear directional expectation, whereas linear models are better suited to more distant applications (e.g., novel tasks or populations). Accordingly, we tested the signatures in Study 6 using i) binary prediction tests, expecting the signatures to selectively predict the attributional statements involving mentalizing, consistent with the original study^84^. We followed the same principles for predictions as outlined above (validation in independent tests), where pattern expression values and predictions are subjected to ROC analyses and binomial tests. In Study 7, we instead used ii) linear-effects models, submitting pattern expression values to repeated-measures ANOVAs across the three social-feedback conditions, followed by Bonferroni-corrected t-tests for pairwise comparisons.

### Region-of-Interest (ROI) Analyses

To test the ability of local patterns to predict mentalizing and to separate self-related versus other-related mentalizing, we trained and cross-validated ROI classifiers using a parallel approach to the whole-brain classifiers. We downloaded a term-based meta-analytic map for ‘Mentalizing’ from NeuroSynth^47^ on 14/09/2022 that included 151 studies. We selected clusters that contained more than 200 voxels, resulting in the following ten ROIs: mPFC, bilateral TPJ, bilateral anterior MTG, precuneus/PCC, right SMA, and three clusters in the cerebellum. In each ROI, we trained and 10-fold cross-validated four classifiers in the training dataset and validated them in the remaining datasets, to (i) predict Self-mentalizing (versus the two remaining conditions), (ii) predict Other-mentalizing (versus the two remaining conditions), (iii) predict mentalizing (Self- and Other-mentalizing versus Control condition), and (iv) differentiate Self-versus Other-processing. We tested these ROI classifiers in the testing datasets using the same parameters and analytic approach as outlined above in the main whole-brain analyses.

### Leave-One-Site-Out classifiers trained on all datasets

To fully benchmark how well our approach generalizes across independent datasets, we trained additional classifiers based on the pooled cohorts from all datasets that used a variation of the trait evaluation task (Studies 1 to 5; *n*=232). We defined five folds, each representing one study, so that the classifiers were trained on four studies and tested on the remaining study in each fold. The cross-validated training of bootstrapped classifiers followed the same analytic flow as the main classifiers (see cross-validated training above). Besides the number of folds, the only other change was the number of bootstrapped samples, which we limited to 2500 samples for computational efficiency. We additionally tested these classifiers in the inference task from Study 6 using the binary prediction tests as described above for the main signatures.

### General statistical approach

All data analysis and data visualizations were performed using Matlab® R2022b software and the Canlab toolbox (https://github.com/canlab). Violin plots in Fig.4 and Fig. 7 were generated using the MATLAB function daviolinplot^91^. Scatter plots in Fig. 5 were generated in R Statistical Software^92^ (v12.1) using ggplot2^93^. Statistical inference used a significance threshold of *p* < 0.05, unless otherwise noted.

## Data availability statement

Source data for all figures and single-subject contrast images of the training dataset (Study 1) are available at https://figshare.com/s/86b77a31fec7cd0e0925. Contrast images from Study 4a can be accessed via OpenNeuro (https://openneuro.org/datasets/ds001618/versions/1.0.1) and data from Study 6 are available through the NIHM Archive (https://nda.nih.gov/edit_collection.html?id=2643). Deidentified data from Studies 2, 3, 4b, 5, and 7 can be made available in line with applicable data protection and privacy regulations upon reasonable request to the corresponding authors.

## Code availability statement

Mentalizing signatures for use in future studies, as well as custom MATLAB scripts for analyses are available at: https://github.com/ldmk/2025_MentalizingSignatures

## Author Contributions

LK, TDW, JAH, and MLS conceived the project. DA and LK conceptualized the research goals, built the collaborations, analyzed the data, and prepared the first draft of the manuscript. DA created the figures. LK, JAH, DAS, AMC, MvB, LK, LZ, LvdM, PFC, EPC, RS, MD, PVr, AT, and PVu designed the experiments and provided data. LK, DA, and LOW acquired funding for the project. LK supervised the project. All authors contributed to the writing and editing of the paper.

## References

1. Moore, C. & Frye, D. The Acquisition and Utility of Theories of Mind. in Children’s Theories of Mind: Mental States and Social Understanding (eds Frye, D. & Moore, C.) 1–14 (Lawrence Erlbaum, Hillsdale, NJ, 1991).

2. Wellman, H. M. Making Minds: How Theory of Mind Develops. (Oxford University Press, 2014).

3. Brüne, M. & Brüne-Cohrs, U. Theory of mind—evolution, ontogeny, brain mechanisms and psychopathology. Neurosci. Biobehav. Rev. 30, 437–455 (2006).

4. Debbané, M. et al. Attachment, Neurobiology, and Mentalizing along the Psychosis Continuum. Front. Hum. Neurosci. 10, (2016).

5. Gray, K., Jenkins, A. C., Heberlein, A. S. & Wegner, D. M. Distortions of mind perception in psychopathology. Proc. Natl. Acad. Sci. 108, 477–479 (2011).

6. Luyten, P., Campbell, C., Allison, E. & Fonagy, P. The Mentalizing Approach to Psychopathology: State of the Art and Future Directions. Annu. Rev. Clin. Psychol. 16, 297–325 (2020).

7. Sharp, C. Mentalizing Problems in Childhood Disorders. in Handbook of Mentalization-Based Treatment 101–121 (John Wiley & Sons, Ltd, 2006). doi:10.1002/9780470712986.ch4.

8. Sloover, M., van Est, L. A. C., Janssen, P. G. J., Hilbink, M. & van Ee, E. A meta-analysis of mentalizing in anxiety disorders, obsessive-compulsive and related disorders, and trauma and stressor related disorders. J. Anxiety Disord. 92, 102641 (2022).

9. Johnson, B. N., Kivity, Y., Rosenstein, L. K., LeBreton, J. M. & Levy, K. N. The association between mentalizing and psychopathology: A meta-analysis of the reading the mind in the eyes task across psychiatric disorders. Clin. Psychol. Sci. Pract. 29, 423–439 (2022).

10. Frith, C. D. & Frith, U. The Neural Basis of Mentalizing. Neuron 50, 531–534 (2006).

11. Saxe, R. Uniquely human social cognition. Curr. Opin. Neurobiol. 16, 235–239 (2006).

12. Schurz, M. et al. Toward a hierarchical model of social cognition: A neuroimaging meta-analysis and integrative review of empathy and theory of mind. Psychol. Bull. 147, 293–327 (2021).

13. Van Overwalle, F. & Baetens, K. Understanding others’ actions and goals by mirror and mentalizing systems: A meta-analysis. NeuroImage 48, 564–584 (2009).

14. Yang, D. Y.-J., Rosenblau, G., Keifer, C. & Pelphrey, K. A. An integrative neural model of social perception, action observation, and theory of mind. Neurosci. Biobehav. Rev. 51, 263–275 (2015).

15. Kragel, P. A., Koban, L., Barrett, L. F. & Wager, T. D. Representation, Pattern Information, and Brain Signatures: From Neurons to Neuroimaging. Neuron 99, 257–273 (2018).

16. Rosenberg, M. D., Casey, B. J. & Holmes, A. J. Prediction complements explanation in understanding the developing brain. Nat. Commun. 9, 589 (2018).

17. Wager, T. D. et al. An fMRI-Based Neurologic Signature of Physical Pain. N. Engl. J. Med. 368, 1388–1397 (2013).

18. Koban, L., Wager, T. D. & Kober, H. A neuromarker for drug and food craving distinguishes drug users from non-users. Nat. Neurosci. 26, 316–325 (2023).

19. Rosenberg, M. D. et al. A neuromarker of sustained attention from whole-brain functional connectivity. Nat. Neurosci. 19, 165–171 (2016).

20. Kim, J. et al. A dorsomedial prefrontal cortex-based dynamic functional connectivity model of rumination. Nat. Commun. 14, 3540 (2023).

21. López-Solà, M. et al. Towards a neurophysiological signature for fibromyalgia. Pain 158, 34–47 (2017).

22. Gabrieli, J. D. E., Ghosh, S. S. & Whitfield-Gabrieli, S. Prediction as a Humanitarian and Pragmatic Contribution from Human Cognitive Neuroscience. Neuron 85, 11–26 (2015).

23. Koban, L. et al. An fMRI-Based Brain Marker of Individual Differences in Delay Discounting. J. Neurosci. 43, 1600–1613 (2023).

24. Woo, C. W., Chang, L. J., Lindquist, M. A. & Wager, T. D. Building better biomarkers: brain models in translational neuroimaging. Nat. Neurosci. 20, 365 (2017).

25. Tamir, D. I. & Thornton, M. A. Modeling the Predictive Social Mind. Trends Cogn. Sci. 22, 201–212 (2018).

26. Qin, P., Wang, M. & Northoff, G. Linking bodily, environmental and mental states in the self—A three-level model based on a meta-analysis. Neurosci. Biobehav. Rev. 115, 77–95 (2020).

27. Corradi-Dell’Acqua, C., Hofstetter, C. & Vuilleumier, P. Cognitive and affective theory of mind share the same local patterns of activity in posterior temporal but not medial prefrontal cortex. Soc. Cogn. Affect. Neurosci. 9, 1175–1184 (2014).

28. Quesque, F. et al. Defining key concepts for mental state attribution. Commun. Psychol. 2, 1–5 (2024).

29. Tan, K. M. et al. Electrocorticographic evidence of a common neurocognitive sequence for mentalizing about the self and others. Nat. Commun. 13, 1919 (2022).

30. Wang, Y. et al. A large-scale structural and functional connectome of social mentalizing. NeuroImage 236, 118115 (2021).

31. Denny, B. T., Kober, H., Wager, T. D. & Ochsner, K. N. A meta-analysis of functional neuroimaging studies of self- and other judgments reveals a spatial gradient for mentalizing in medial prefrontal cortex. J. Cogn. Neurosci. 24, 1742–1752 (2012).

32. Parelman, J. M. et al. Overlapping Functional Representations of Self- and Other-Related Thought are Separable Through Multivoxel Pattern Classification. Cereb. Cortex 10.1093/CERCOR/BHAB272 (2021) doi:10.1093/CERCOR/BHAB272.

33. Fossati, P. et al. In Search of the Emotional Self: An fMRI Study Using Positive and Negative Emotional Words. Am. J. Psychiatry 160, 1938–1945 (2003).

34. Murray, R. J., Debbané, M., Fox, P. T., Bzdok, D. & Eickhoff, S. B. Functional connectivity mapping of regions associated with self- and other-processing. Hum. Brain Mapp. 36, 1304–1324 (2014).

35. Northoff, G. et al. Self-referential processing in our brain—A meta-analysis of imaging studies on the self. NeuroImage 31, 440–457 (2006).

36. Van Der Meer, L., Costafreda, S., Aleman, A. & David, A. S. Self-reflection and the brain: A theoretical review and meta-analysis of neuroimaging studies with implications for schizophrenia. Neurosci. Biobehav. Rev. 34, 935–946 (2010).

37. Arioli, M., Cattaneo, Z., Parimbelli, S. & Canessa, N. Relational vs representational social cognitive processing: a coordinate-based meta-analysis of neuroimaging data. Soc. Cogn. Affect. Neurosci. 18, nsad003 (2023).

38. Tamir, D. I., Bricker, A. B., Dodell-Feder, D. & Mitchell, J. P. Reading fiction and reading minds: the role of simulation in the default network. Soc. Cogn. Affect. Neurosci. 11, 215–224 (2016).

39. Wagner, D. D., Chavez, R. S. & Broom, T. W. Decoding the neural representation of self and person knowledge with multivariate pattern analysis and data-driven approaches. Wiley Interdiscip. Rev. Cogn. Sci. 10, (2019).

40. Arioli, M., Cattaneo, Z., Ricciardi, E. & Canessa, N. Overlapping and specific neural correlates for empathizing, affective mentalizing, and cognitive mentalizing: A coordinate-based meta-analytic study. Hum. Brain Mapp. 42, 4777–4804 (2021).

41. Molapour, T. et al. Seven computations of the social brain. Soc. Cogn. Affect. Neurosci. 16, 745–760 (2021).

42. Kelley, W. M. et al. Finding the Self? An Event-Related fMRI Study. J. Cogn. Neurosci. 14, 785–794 (2002).

43. Koban, L., Pichon, S. & Vuilleumier, P. Responses of medial and ventrolateral prefrontal cortex to interpersonal conflict for resources. Soc. Cogn. Affect. Neurosci. 9, 561–569 (2014).

44. Scheinost, D. et al. Ten simple rules for predictive modeling of individual differences in neuroimaging. NeuroImage 193, 35–45 (2019).

45. Bora, E., Yucel, M. & Pantelis, C. Theory of mind impairment in schizophrenia: Meta-analysis. Schizophr. Res. 109, 1–9 (2009).

46. Sprong, M., Schothorst, P., Vos, E., Hox, J. & Engeland, H. V. Theory of mind in schizophrenia: Meta-analysis. Br. J. Psychiatry 191, 5–13 (2007).

47. Yarkoni, T., Poldrack, R. A., Nichols, T. E., Van Essen, D. C. & Wager, T. D. Large-scale automated synthesis of human functional neuroimaging data. Nat. Methods 8, 665–670 (2011).

48. Oosterwijk, S., Snoek, L., Rotteveel, M., Barrett, L. F. & Scholte, H. S. Shared states: using MVPA to test neural overlap between self-focused emotion imagery and other-focused emotion understanding. Soc. Cogn. Affect. Neurosci. 12, 1025–1035 (2017).

49. Van Overwalle, F. & Vandekerckhove, M. Implicit and explicit social mentalizing: dual processes driven by a shared neural network. Front. Hum. Neurosci. 7, (2013).

50. Heatherton, T. F. et al. Medial prefrontal activity differentiates self from close others. Soc. Cogn. Affect. Neurosci. 1, 18–25 (2006).

51. Koban, L., Gianaros, P. J., Kober, H. & Wager, T. D. The self in context: brain systems linking mental and physical health. Nat. Rev. Neurosci. 10.1038/s41583-021-00446-8 (2021) doi:10.1038/s41583-021-00446-8.

52. Chavez, R. S., Heatherton, T. F. & Wagner, D. D. Neural Population Decoding Reveals the Intrinsic Positivity of the Self. Cereb. Cortex cercor;bhw302v1 (2017) doi:10.1093/cercor/bhw302.

53. Northoff, G. & Hayes, D. J. Is Our Self Nothing but Reward? Biol. Psychiatry 69, 1019–1025 (2011).

54. Tamir, D. I. & Mitchell, J. P. Disclosing information about the self is intrinsically rewarding. Proc. Natl. Acad. Sci. 109, 8038–8043 (2012).

55. Maresh, E. L. & Andrews-Hanna, J. R. Putting the “Me” in “Mentalizing”: Multiple Constructs Describing Self Versus Other During Mentalizing and Implications for Social Anxiety Disorder. in The Neural Basis of Mentalizing (eds Gilead, M. & Ochsner, K. N.) 629–658 (Springer International Publishing, Cham, 2021). doi:10.1007/978-3-030-51890-5_33.

56. Babo-Rebelo, M. & Tallon-Baudry, C. Interoceptive signals, brain dynamics, and subjectivity. in The Interoceptive Mind: From Homeostasis to Awareness (eds Tsakiris, M. & De Preester, H.) 46–62 (Oxford University Press, Oxford, UK, 2018).

57. Garfinkel, S. N., Nagai, Y., Seth, A. K. & Critchley, H. D. Neuroimaging Studies of Interoception and Self-Awareness. in Neuroimaging of Consciousness (eds Cavanna, A. E., Nani, A., Blumenfeld, H. & Laureys, S.) 207–224 (Springer, Berlin, Heidelberg, 2013). doi:10.1007/978-3-642-37580-4_11.

58. Andrews-Hanna, J. R., Reidler, J. S., Sepulcre, J., Poulin, R. & Buckner, R. L. Functional-Anatomic Fractionation of the Brain’s Default Network. Neuron 65, 550–562 (2010).

59. Peer, M., Salomon, R., Goldberg, I., Blanke, O. & Arzy, S. Brain system for mental orientation in space, time, and person. Proc. Natl. Acad. Sci. 112, 11072–11077 (2015).

60. Schurz, M., Radua, J., Aichhorn, M., Richlan, F. & Perner, J. Fractionating theory of mind: A meta-analysis of functional brain imaging studies. Neurosci. Biobehav. Rev. 42, 9–34 (2014).

61. Potvin, S., Gamache, L. & Lungu, O. A Functional Neuroimaging Meta-Analysis of Self-Related Processing in Schizophrenia. Front. Neurol. 10, (2019).

62. Crone, E. A. & Fuligni, A. J. Self and Others in Adolescence. Annu. Rev. Psychol. 71, 447–469 (2020).

63. Fehlbaum, L. V., Borbás, R., Paul, K., Eickhoff, S. B. & Raschle, N. M. Early and late neural correlates of mentalizing: ALE meta-analyses in adults, children and adolescents. Soc. Cogn. Affect. Neurosci. 17, 351–366 (2022).

64. Spunt, R. P. & Adolphs, R. Folk Explanations of Behavior: A Specialized Use of a Domain-General Mechanism. Psychol. Sci. 26, 724–736 (2015).

65. Spunt, R. P. & Adolphs, R. A new look at domain specificity: insights from social neuroscience. Nat. Rev. Neurosci. 18, 559–567 (2017).

66. Binney, R. J. & Ramsey, R. Social Semantics: The role of conceptual knowledge and cognitive control in a neurobiological model of the social brain. Neurosci. Biobehav. Rev. 112, 28–38 (2020).

67. Andrews-Hanna, J. R. & Grilli, M. D. Mapping the Imaginative Mind: Charting New Paths Forward. Curr. Dir. Psychol. Sci. 30, 82–89 (2021).

68. Baetens, K., Ma, N., Steen, J. & Van Overwalle, F. Involvement of the mentalizing network in social and non-social high construal. Soc. Cogn. Affect. Neurosci. 9, 817–824 (2014).

69. Andrews-Hanna, J. R., Smallwood, J. & Spreng, R. N. The default network and self-generated thought: component processes, dynamic control, and clinical relevance. Ann. N. Y. Acad. Sci. 1316, 29–52 (2014).

70. Yeshurun, Y., Nguyen, M. & Hasson, U. The default mode network: where the idiosyncratic self meets the shared social world. Nat. Rev. Neurosci. 22, 181–192 (2021).

71. Defendini, A. & Jenkins, A. C. Dissociating neural sensitivity to target identity and mental state content type during inferences about other minds. Soc. Neurosci. 18, 103–121 (2023).

72. Kim, H. J., Lux, B. K., Lee, E., Finn, E. S. & Woo, C.-W. Brain decoding of spontaneous thought: Predictive modeling of self-relevance and valence using personal narratives. Proc. Natl. Acad. Sci. 121, e2401959121 (2024).

73. Kim Lux, B., Andrews-Hanna, J. R., Han, J., Lee, E. & Woo, C.-W. When self comes to a wandering mind: Brain representations and dynamics of self-generated concepts in spontaneous thought. Sci. Adv. 8, eabn8616 (2022).

74. Courtney, A. L. & Meyer, M. L. Self-Other Representation in the Social Brain Reflects Social Connection. J. Neurosci. 40, 5616–5627 (2020).

75. Debbané, M. et al. Brain activity underlying negative self- and other-perception in adolescents: The role of attachment-derived self-representations. Cogn. Affect. Behav. Neurosci. 17, 554–576 (2017).

76. van Buuren, M. et al. Neural correlates of self- and other-referential processing in young adolescents and the effects of testosterone and peer similarity. NeuroImage 219, 117060 (2020).

77. Fuentes-Claramonte, P. et al. Shared and differential default-mode related patterns of activity in an autobiographical, a self-referential and an attentional task. PLOS ONE 14, e0209376 (2019).

78. Fuentes-Claramonte, P. et al. Brain imaging correlates of self- and other-reflection in schizophrenia. NeuroImage Clin. 25, 102134 (2020).

79. First, M. B. Structured Clinical Interview for the DSM (SCID). in The Encyclopedia of Clinical Psychology 1–6 (John Wiley & Sons, Ltd, 2015). doi:10.1002/9781118625392.wbecp351.

80. Zhang, L., Opmeer, E. M., Ruhé, H. G., Aleman, A. & van der Meer, L. Brain activation during self- and other-reflection in bipolar disorder with a history of psychosis: Comparison to schizophrenia. NeuroImage Clin. 8, 202–209 (2015).

81. Sheehan, D. V. et al. The Mini-International Neuropsychiatric Interview (M.I.N.I.): the development and validation of a structured diagnostic psychiatric interview for DSM-IV and ICD-10. J. Clin. Psychiatry 59 Suppl 20, 22-33;quiz 34-57 (1998).

82. Kemp, R. & David, A. Insight and compliance. in Treatment compliance and the therapeutic alliance (ed. Blackwell, B.) 61–84 (Harwood Academic Publishers, 1997).

83. Beck, A. T., Baruch, E., Balter, J. M., Steer, R. A. & Warman, D. M. A new instrument for measuring insight: the Beck Cognitive Insight Scale. Schizophr. Res. 68, 319–329 (2004).

84. Tusche, A., Spunt, R. P., Paul, L. K., Tyszka, J. M. & Adolphs, R. Neural signatures of social inferences predict the number of real-life social contacts and autism severity. Nat. Commun. 14, 4399 (2023).

85. Ma, S. et al. When Empathy Gets Tough: Neural Responses to Overcoming the Self in a Novel Paradigm Predict Everyday Prosocial Behavior. (2024) doi:10.31234/osf.io/qcj45.

86. Cox, D. D. & Savoy, R. L. Functional magnetic resonance imaging (fMRI) “brain reading”: detecting and classifying distributed patterns of fMRI activity in human visual cortex. NeuroImage 19, 261–270 (2003).

87. Misaki, M., Kim, Y., Bandettini, P. A. & Kriegeskorte, N. Comparison of multivariate classifiers and response normalizations for pattern-information fMRI. NeuroImage 53, 103–118 (2010).

88. Mourão-Miranda, J., Bokde, A. L. W., Born, C., Hampel, H. & Stetter, M. Classifying brain states and determining the discriminating activation patterns: Support Vector Machine on functional MRI data. NeuroImage 28, 980–995 (2005).

89. Woo, C.-W. et al. Separate neural representations for physical pain and social rejection. Nat. Commun. 5, 5380 (2014).

90. Steardo, L. et al. Application of Support Vector Machine on fMRI Data as Biomarkers in Schizophrenia Diagnosis: A Systematic Review. Front. Psychiatry 11, 588 (2020).

91. Karvelis, P. violin and raincloud plots (https://github.com/povilaskarvelis/DataViz), GitHub.

92. R Core Team. R: A language and environment for statistical computing. R Foundation for Statistical Computing, Vienna, Austria. (2024).

93. Wickham, H. ggplot2: Elegant Graphics for Data Analysis. Springer-Verlag. (2016).

