## Supplementary for "Brain neuromarkers predict self- and other-related mentalizing across adult, clinical, and developmental samples"

### Supplementary Table 1

*Demographics*

| Study | Population | n | Gender (females) | Age mean (SD) | Handedness (left) | Education (in years) | Place | References |
|---|---|---|---|---|---|---|---|---|
| Study 1 | Healthy adults | 21 | 10 | 23.7 (7.4) | 2 | N/A | Switzerland | Koban et al., 2014 |
| Study 2 | Adolescents | 44 | 23 | 16.0 (1.9) | 0 | N/A | Switzerland | Debbane et al., 2017 |
| Study 3 | Adolescents | 61 | 27 | 12.9 (0.4) | 11 | N/A | The Netherlands | van Buuren et al., 2020 |
| Study 4a | Healthy adults | 33 | 14 | 41.7 (11.7) | 0 | 12.7 | Spain | Fuentes-Claramonte et al., 2019 & 2020 |
| Study 4b | Schizophrenia patients | 23 | 7 | 37.0 (8.1) | 0 | 10.4 | | |
| Study 5a | Healthy adults | 15 | 6 | 33.3 (11.3) | 2 | 17.2 | The Netherlands | Zhang et al., 2015 |
| Study 5b | Schizophrenia patients | 17 | 6 | 35.5 (9.7) | 1 | 16.6 | | |
| Study 5c | Bipolar patients | 18 | 9 | 40.3 (12.7) | 0 | 17.0 | | |
| Study 6a | Healthy adults | 59 | 26 | 28.3 (5.2) | 2 | N/A | USA | Tusche et al., 2023 |
| Study 6b | Healthy adults | 50 | 19 | 33.6 (7.3) | 4 | N/A | USA | Tusche et al., 2023 |
| Study 7 | Healthy adults | 49 | 26 | 22.7 (3.9) | 6 | 13 | USA | Ma et al., 2024 |
| **Overall** | | **390** | **173** | **26.9 (11.7)** | **28** | | | |

*Note.* NA = not assessed or inaccessible. Links to the published studies are as follows: Study 2 (doi.org/10.3758/s13415-017-0497-9); Study 3 (doi.org/10.1016/j.neuroimage.2020.117060); Study 4, 2019 (doi.org/10.1371/journal.pone.0209376); Study 4, 2020 (doi.org/10.1016/j.nicl.2019.102134); Study 5 (doi.org/10.1016/j.nicl.2015.04.010); Study 6 (doi.org/10.1038/s41467-023-40078-3); Study 7 (doi.org/10.31234/osf.io/qcj45)

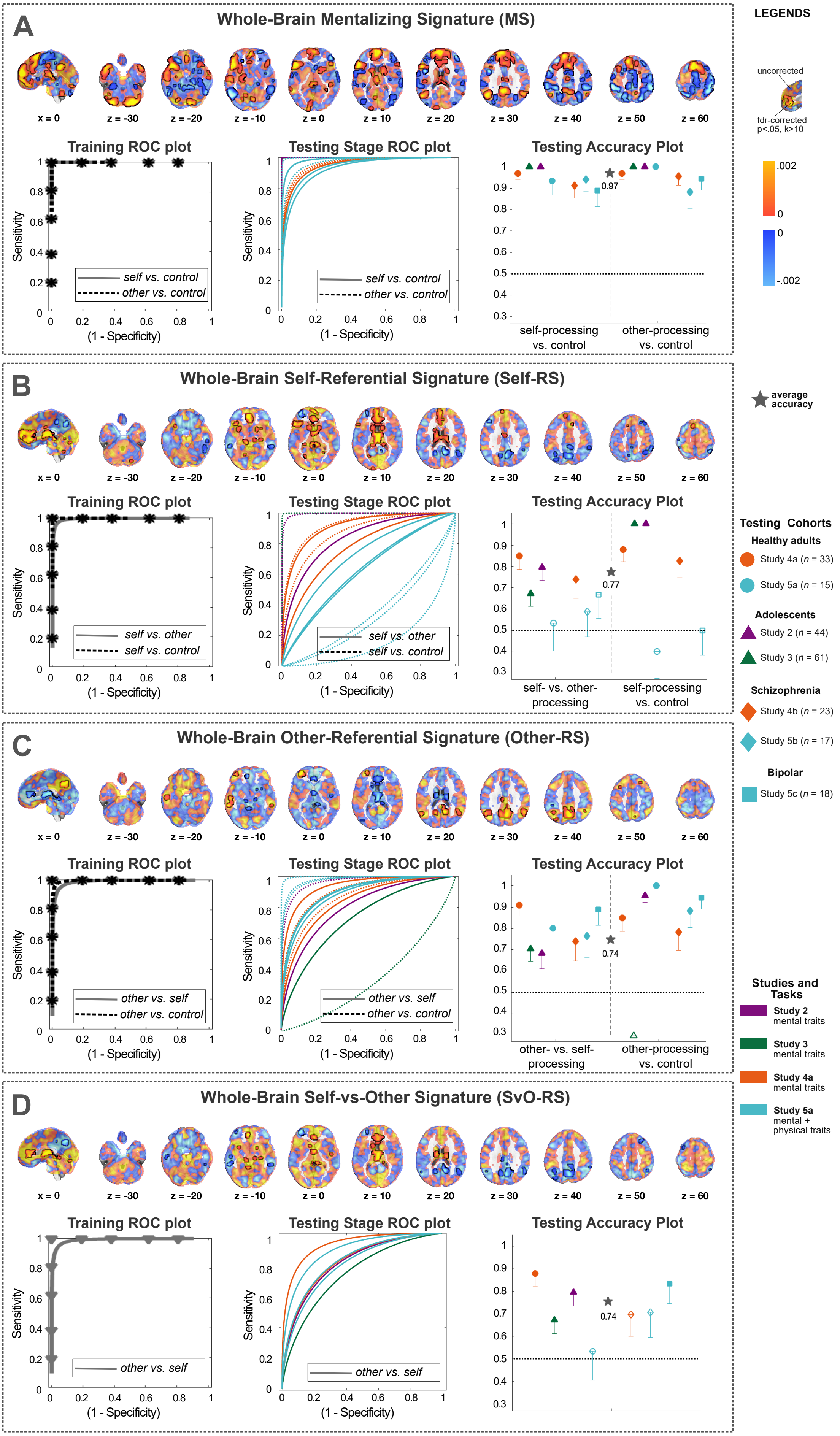
A
Whole-Brain Mentalizing Signature (MS)
x = 0
z = -30
z = -20
z = -10
z = 0
z = 10
z = 20
z = 30
z = 40
z = 50
z = 60
Training ROC plot
Sensitivity
(1 - Specificity)
self vs. control
other vs. control
Testing Stage ROC plot
Testing Accuracy Plot
0.97
self-processing vs. control
other-processing vs. control
B
Whole-Brain Self-Referential Signature (Self-RS)
self vs. other
self vs. control
0.77
self- vs. other-processing
self-processing vs. control
C
Whole-Brain Other-Referential Signature (Other-RS)
other vs. self
other vs. control
0.74
other- vs. self-processing
other-processing vs. control
D
Whole-Brain Self-vs-Other Signature (SvO-RS)
other vs. self
0.74
LEGENDS
uncorrected
fdr-corrected p<.05, k>10
.002
0
0
-.002
average accuracy
Testing Cohorts
Healthy adults
Study 4a (n = 33)
Study 5a (n = 15)
Adolescents
Study 2 (n = 44)
Study 3 (n = 61)
Schizophrenia
Study 4b (n = 23)
Study 5b (n = 17)
Bipolar
Study 5c (n = 18)
Studies and Tasks
Study 2 mental traits
Study 3 mental traits
Study 4a mental traits
Study 5a mental + physical traits

**Supplementary Figure 1 — Whole Brain Classifiers (n=21).** In order to determine whether the use of a social-cognition mask may have altered the results substantially, we trained new classifiers using the same analytic flow (n=21) but using whole-brain contrast images (with a grey-matter mask). On the top row of each box, weight maps illustrate the positive and negative weights for each classifier. Voxels significant at an FDR-corrected threshold ($q < .05$, minimum cluster size of $k = 10$) are highlighted by black outlines. The left and middle plots in each box depict the receiver operating characteristic (ROC) plots from the i) cross-validated training and ii) testing datasets. The last column shows the accuracies of all three mentalizing signatures in each testing dataset separately, with color representing the study, and shape illustrating the cohort type. The star shows the weighted average accuracy across all seven datasets. These additional analyses illustrate that masking the contrast images used in this study did not alter the results substantially, as the whole-brain approach resulted with comparable prediction accuracies and brain weights.

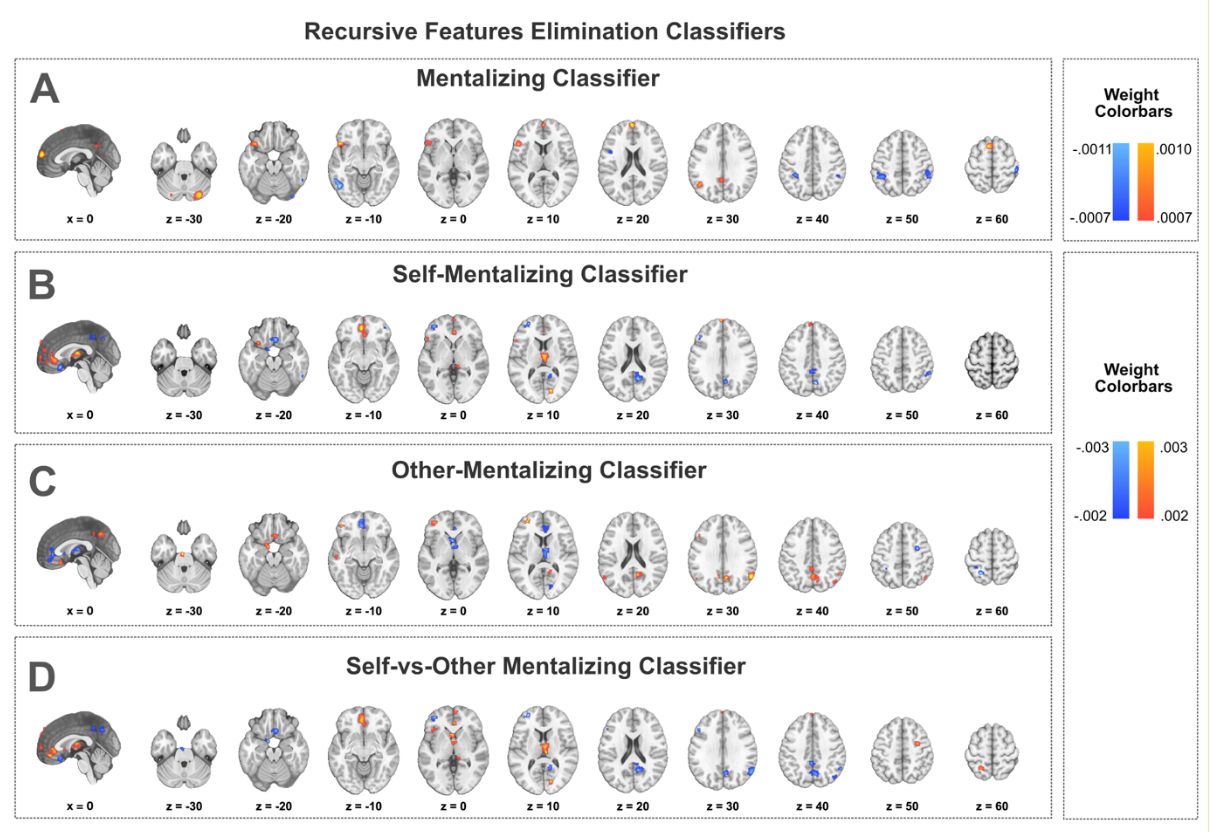


**Supplementary Figure 2 — Recursive Feature Elimination (RFE) Classifiers (n=21**). Additional classifiers using a recursive feature elimination method yielded similar results as the original bootstrapping approach. Linear SVM models with 10-fold cross-validation were trained in multiple iterations, in which 5000 voxels were removed from the model in each step to reach the final voxel size of 2000. The final models with 2000 features are illustrated here, including only clusters with a minimum voxel size of 20. All four final models had 100% prediction accuracies in the cross-validated training phase.

### Supplementary Table 2

*Significant clusters of the Self-RS*

| Labeled Name | Atlas region (see note) | x | y | z | Voxel number | Max(z) |
|---|---|---|---|---|---|---|
| Cerebellum | Cblm Crus II R | 26 | -80 | -38 | 54 | 0.00035 |
| Ventral Anterior Cingulate Cortex (vACC) / ventromedial prefrontal cortex (vmPFC) (incl. dorsomedial and frontopolar prefrontal cortices) | Multiple regions | -2 | 44 | 0 | 933 | 0.00068 |
| Anterior insula (AI) | AAIC L | -34 | 10 | -10 | 33 | 0.00043 |
| Caudate nucleus | Caudate Ca R | 18 | 10 | -6 | 13 | 0.00033 |
| Inferior frontal gyrus (IFG) | 45 L | -46 | 24 | 2 | 162 | 0.00057 |
| Thalamus | Multiple regions | 0 | -12 | 10 | 311 | 0.00068 |
| Caudate nucleus | Caudate Ca R | 12 | 14 | 8 | 17 | 0.00031 |
| Caudate nucleus | Caudate Ca L | -14 | 14 | 6 | 32 | 0.00035 |
| Dorsolateral prefrontal cortex (dlPFC) | 9m L | -8 | 48 | 38 | 21 | 0.00037 |
| Superior frontal lobule/ dorsomedial prefrontal cortex (dmPFC) | SFL L | -8 | 22 | 56 | 46 | 0.00040 |
| Temporal pole | TGd L | -34 | 14 | -34 | 13 | -0.00050 |
| Inferior temporal gyrus/sulcus | TE2p R | 46 | -46 | -22 | 27 | -0.00045 |
| Posterior temporal cortex | PH R | 52 | -54 | -14 | 21 | -0.00054 |
| dlPFC | a9 46v L | -38 | 48 | 4 | 26 | -0.00072 |
| Posterior cingulate cortex (PCC) | POS2 R | 6 | -60 | 32 | 270 | -0.00070 |
| Inferior parietal cortex (IPC) / Temporoparietal junction (TPJ) | PGi R | 52 | -62 | 22 | 52 | -0.00046 |
| IPC / TPJ | PFm R | 52 | -50 | 24 | 20 | -0.00035 |
| IPC / TPJ | PGs L | -34 | -76 | 40 | 14 | -0.00044 |
| PCC | 31a L | -2 | -42 | 42 | 20 | -0.00054 |
| IPC / TPJ | PFm R | 52 | -44 | 48 | 28 | -0.00064 |

*Note.* Significant positive and negative weights contributing to the Self-Referential Signature (Self-RS). FDR-corrected $p < 0.05$ with cluster size $k > 10$ across the masked whole-brain. Cortical atlas regions are labeled based on a combination of parcellations available on GitHub: https://github.com/canlab/Neuroimaging_Pattern_Masks/tree/master/Atlases_and_parcellations/2018_Wager_combined_atlas. L refers to the left hemisphere, and R refers to the right hemisphere.

**Supplementary Table 3**

*Significant clusters of the Other-RS*

| Labeled Name | Atlas region (see note) | x | y | z | Voxel number | Max(z) |
|---|---|---|---|---|---|---|
| Superior temporal sulcus (STS) | STSda L | -56 | -16 | -12 | 174 | 0.00053 |
| dlPFC | a9 46v L | -38 | 50 | 2 | 90 | 0.00085 |
| PCC/Precuneus | Multiple regions | 4 | -60 | 34 | 733 | 0.00081 |
| IPC / TPJ | PGi L | -52 | -60 | 26 | 245 | 0.00048 |
| IPC / TPJ | PGi R | 52 | -58 | 30 | 339 | 0.00058 |
| Retrosplenial cortex | RSC L | -4 | -28 | 28 | 15 | 0.00039 |
| IPC / TPJ | PGs L | -36 | -76 | 42 | 14 | 0.00048 |
| IPC / TPJ | PFm L | -42 | -58 | 42 | 38 | 0.00034 |
| Anterior insula (AI) | AAIC R | 30 | 12 | -10 | 15 | -0.00034 |
| vACC/vmPFC | Multiple regions | -2 | 40 | 2 | 498 | -0.00058 |
| Caudate nucleus | Caudate Ca R | 20 | 8 | -6 | 20 | -0.00032 |
| Thalamus | Bstem Midbd R | 12 | -30 | -2 | 24 | -0.00049 |
| Anterior insula (AI) | AAIC L | -34 | 10 | -6 | 13 | -0.00027 |
| Ventral striatum | V Striatum L | -2 | 16 | -2 | 13 | -0.00045 |
| Thalamus | Multiple regions | 0 | -10 | 10 | 314 | -0.00059 |
| Caudate nucleus | Caudate Ca L | -12 | 14 | 6 | 24 | -0.00036 |
| Ventral striatum | Cau R | 0 | 24 | 8 | 13 | -0.00035 |
| Middle temporal complex (MT+) | LO3 L | -44 | -80 | 12 | 17 | -0.00026 |
| IPC | IP2 L | -46 | -40 | 42 | 10 | -0.00036 |
| Somatosensory cortex | 2 L | -30 | -44 | 54 | 19 | -0.00033 |

*Note.* Significant positive and negative weights contributing to the Other-Referential Signature (MS-Other). FDR-corrected $p < 0.05$ with cluster size $k > 10$ across the masked whole-brain. Cortical atlas regions are labeled based on a combination of parcellations available on GitHub: https://github.com/canlab/Neuroimaging_Pattern_Masks/tree/master/Atlases_and_parcellations/2018_Wager_combined_atlas. L refers to the left hemisphere, and R refers to the right hemisphere.

**Supplementary Table 4**

*Significant clusters of the MS*

| Labeled Name | Atlas region (see note) | x | y | z | Voxel number | Max(z) |
|---|---|---|---|---|---|---|
| Cerebellum | Cblm IX R | 2 | -54 | -44 | 109 | 0.00016 |
| Cerebellum | Cblm CrusI R | 28 | -82 | -32 | 597 | 0.00040 |
| Cerebellum | Cblm CrusI L | -24 | -80 | -34 | 373 | 0.00028 |
| STS/IFG (incl. vlPFC, AI, STG, temporal pole) | Multiple regions | -48 | 18 | -10 | 2536 | 0.00038 |
| Temporal pole | TGd R | 50 | 16 | -30 | 114 | 0.00014 |
| Orbitofrontal cortex (OFC) | 10v R | 0 | 38 | -20 | 15 | 0.00012 |
| IFG/ventrolateral PFC (vlPFC) | 45 R | 44 | 24 | -10 | 195 | 0.00014 |
| Amygdala | Bstem Ponscd | -24 | -12 | -10 | 68 | 0.00010 |
| Dorsal anterior cingulate cortex (dACC)/vACC (incl. dmPFC, vmPFC, dlPFC, frontal eye fields) | Multiple regions | -6 | 42 | 36 | 3532 | 0.00036 |
| STS | STSdp R | 46 | -28 | 0 | 20 | 0.00010 |
| Putamen | Putamen Pa R | 30 | 12 | 0 | 13 | 0.00007 |
| Thalamus | Bstem_SC R | 4 | -28 | 2 | 10 | 0.00015 |
| Caudate nucleus | Caudate Ca R | 16 | 10 | 10 | 107 | 0.00012 |
| Thalamus | Thal MD | -2 | -16 | 10 | 96 | 0.00025 |
| IPC / TPJ | PGi L | -48 | -62 | 26 | 512 | 0.00023 |
| Posterior opercular cortex | OP1 L | -50 | -24 | 18 | 12 | 0.00008 |
| IPC / TPJ | PGi R | 52 | -58 | 30 | 48 | 0.00012 |
| PCC | Multiple regions | -2 | -50 | 32 | 544 | 0.00023 |
| dlPFC | 9p R | 18 | 40 | 36 | 42 | 0.00010 |
| Supplementary motor area (SMA) | 8Av L | -36 | 12 | 46 | 68 | 0.00012 |
| dlPFC | 8BL R | 14 | 26 | 52 | 19 | 0.00008 |
| Perirhinal ectorhinal cortex | PeEc L | -34 | 0 | -38 | 17 | -0.00010 |
| Amygdala | Amygdala LB | 26 | 0 | -26 | 13 | -0.00012 |
| Cerebellum | Cblm VI L | -36 | -42 | -24 | 21 | -0.00011 |
| Middle temporal gyrus (MTG) posterior | TE1p R | 54 | -52 | -14 | 215 | -0.00024 |
| MTG anterior | TE2a R | 48 | -18 | -22 | 27 | -0.00012 |
| Hippocampus | H R | 20 | -16 | -20 | 29 | -0.00020 |
| Entorhinal cortex/amygdala | Pir R | 36 | 2 | -18 | 29 | -0.00014 |
| Parahippocampal gyrus | PHA3 L | -32 | -32 | -20 | 10 | -0.00010 |
| Amygdala | Amygdala LB | 18 | 4 | -18 | 29 | -0.00015 |
| Posterior temporal cortex/MT+ | PH L | -52 | -58 | -10 | 419 | -0.00029 |
| OFC | 11l R | 20 | 54 | -14 | 34 | -0.00019 |
| MTG medial | TE1m R | 60 | -26 | -16 | 12 | -0.00011 |

| | | | | | | |
|---|---|---|---|---|---|---|
| Posterior temporal cortex | PH R | 50 | -70 | -8 | 43 | -0.00017 |
| Auditory association cortex | A5 R | 60 | -6 | -2 | 54 | -0.00020 |
| STS | STSvp R | 56 | -40 | -4 | 12 | -0.00012 |
| Middle Insula | MI L | -34 | 14 | 0 | 18 | -0.00010 |
| Inferior frontal sulcus | IFSa R | 48 | 38 | 6 | 57 | -0.00016 |
| Extrastriate cortex/MTG | TPOJ2 R | 48 | -64 | 12 | 327 | -0.00013 |
| Premotor cortex/Inferior frontal sulcus | 6r L | -46 | 6 | 26 | 333 | -0.00023 |
| dlPFC | a9 46v L | -36 | 48 | 10 | 17 | -0.00016 |
| Extrastriate cortex | V3CD R | 36 | -84 | 12 | 32 | -0.00011 |
| IPC | PGp L | -44 | -80 | 16 | 38 | -0.00009 |
| Superior temporal gyrus (STG) | STV R | 58 | -42 | 18 | 35 | -0.00012 |
| Supramarginal gyrus/IPC | PF L | -44 | -42 | 44 | 629 | -0.00029 |
| TPJ/Angular gyrus (AG) | PSL R | 60 | -30 | 18 | 18 | -0.00011 |
| dlPFC | p9 46v L | -40 | 30 | 22 | 10 | -0.00010 |
| Premotor cortex | 6v R | 50 | 14 | 28 | 32 | -0.00015 |
| IPC | IP1 R | 30 | -64 | 42 | 151 | -0.00016 |
| IPC/TPJ | PFm R | 46 | -42 | 48 | 576 | -0.00033 |
| PCC | 5mv L | -10 | -38 | 46 | 121 | -0.00019 |
| Superior parietal cortex/medial intraparietal sulcus | MIP L | -24 | -62 | 50 | 75 | -0.00018 |
| MCC | 24dd R | 10 | -16 | 42 | 14 | -0.00006 |
| SMA | SCEF R | 0 | 4 | 52 | 284 | -0.00018 |
| Premotor cortex/SMA | 55b R | 50 | -4 | 44 | 31 | -0.00013 |
| SMA | 6a L | -28 | 0 | 50 | 20 | -0.00010 |
| SMA | 8Av R | 32 | 30 | 50 | 13 | -0.00021 |
| SMA | 6a L | -24 | 8 | 54 | 23 | -0.00012 |
| Primary sensory cortex | 1 R | 46 | -28 | 62 | 35 | -0.00036 |

*Note.* Significant positive and negative weights contributing to the Mentalizing Signature (MS). FDR-corrected $p < 0.05$ with cluster size $k > 10$ across the masked whole-brain. Cortical atlas regions are labeled based on a combination of parcellations available on GitHub: https://github.com/canlab/Neuroimaging_Pattern_Masks/tree/master/Atlases_and_parcellations/2018_Wager_combined_atlas. L refers to the left hemisphere, and R refers to the right hemisphere.

**Supplementary Table 5**

*Significant clusters of the Self-vs-Other Referential Signature (SvO-RS)*

| **Labeled Name** | **Atlas region (see note)** | **x** | **y** | **z** | **Voxel number** | **Max(z)** |
|---|---|---|---|---|---|---|
| dACC, vACC, vmPFC (bilateral) | Multiple regions | -2 | 40 | 0 | 501 | 0.00061 |
| Anterior insula | AAIC L | -34 | 10 | -8 | 17 | 0.00027 |
| IFS | 47l L | -38 | 28 | -2 | 17 | 0.00035 |
| Thalamus (bilateral) | Multiple regions | 0 | -10 | 8 | 271 | 0.00055 |
| Caudate nucleus | Caudate Ca L | -12 | 14 | 6 | 25 | 0.00034 |
| Ventral striatum | V Striatum L | 0 | 24 | 8 | 11 | 0.00036 |
| Middle temporal complex (MT+) | LO3 L | -48 | -78 | 10 | 10 | 0.00023 |
| Superior temporal sulcus (STS) | STSda L | -56 | -16 | -12 | 61 | -0.00045 |
| Inferior frontal cortex | p47r L | -38 | 48 | 2 | 49 | -0.00079 |
| PCC (incl. precuneus) | 7m R | 4 | -60 | 34 | 484 | -0.00073 |
| IPC / TPJ | PGi R | 52 | -60 | 24 | 82 | -0.00045 |
| IPC / TPJ | PFm L | -56 | -58 | 26 | 49 | -0.00044 |
| IPC / TPJ | PGs L | -34 | -76 | 42 | 12 | -0.00044 |
| IPC / TPJ | PFm L | -40 | -58 | 42 | 14 | -0.00035 |

*Note.* Significant positive and negative weights contributing to the Self-vs-Other Referential Signature (SvO-RS). FDR-corrected $p < 0.05$ with cluster size $k > 10$ across the masked whole-brain. Cortical atlas regions are labeled based on a combination of parcellations available on GitHub: https://github.com/canlab/Neuroimaging_Pattern_Masks/tree/master/Atlases_and_parcellations/2018_Wager_combined_atlas. L refers to the left hemisphere, and R refers to the right hemisphere.

### Table S6

*Prediction results of the mentalizing signatures across training and validation datasets*

***Self-Mentalizing Signature (MS-Self)***

| **Dataset** | **Study** | **Sample** | **n** | **Classification Task** | **Prediction Outcome** |
|---|---|---|---|---|---|
| Training | Study 1 | Healthy adult | 21 | Self vs Other | acc.=1.00(+/-0.00), *P*<0.0001***, sens.=1.00, spec.=1.00, AUC=1.00, *d*=2.61 |
| | | | | Self vs Control | acc.=1.00(+/-0.00), *P*<0.0001***, sens.=1.00, spec.=1.00, AUC=1.00, *d*=3.74 |
| Validation | Study 2 | Adolescent | 44 | Self vs Other | acc.=0.77(+/-0.06), P=0.0004***, sens.=0.77, spec.=0.77, AUC=0.88, *d*=1.13 |
| | | | | Self vs Control | acc.=0.98(+/-0.02), *P*<0.0001***, sens.=0.98, spec.=0.98, AUC=1.00, *d*=2.60 |
| Validation | Study 3 | Adolescent | 61 | Self vs Other | acc.=0.75(+/-0.06), *P*<0.0001***, sens.=0.75, spec.=0.75, AUC=0.84, *d*=0.95 |
| | | | | Self vs Control | acc.=1.00(+/-0.00), *P*<0.0001***, sens.=1.00, spec.=1.00, AUC=1.00, *d*=3.52 |
| Validation | Study 4a | Healthy adult | 33 | Self vs Other | acc.=0.91(+/-0.05), *P*<0.0001***, sens.=0.91, spec.=0.91, AUC=0.98, *d*=1.83 |
| | | | | Self vs Control | acc.=0.82(+/-0.07), *P*=0.0003***, sens.=0.82, spec.=0.82, AUC=0.93, *d*=1.51 |
| Validation | Study 4b | Schizophrenia patient | 23 | Self vs Other | acc.=0.78(+/-0.09), *P*=0.0106***, sens.=0.78, spec.=0.78, AUC=0.71, *d*=0.50 |
| | | | | Self vs Control | acc.=0.83(+/-0.08), *P*=0.0026***, sens.=0.83, spec.=0.83, AUC=0.87, *d*=1.01 |
| Validation | Study 5a | Healthy adult | 15 | Self vs Other | acc.=0.47(+/-0.13), *P*>0.20, sens.=0.47, spec.=0.47, AUC=0.71, *d*=0.61 |
| | | | | Self vs Control | acc.=0.53(+/-0.13), *P*>0.20, sens.=0.53, spec.=0.53, AUC=0.60, *d*=0.31 |
| Validation | Study 5b | Schizophrenia patient | 17 | Self vs Other | acc.=0.76(+/-0.10), *P*=0.0490***, sens.=0.76, spec.=0.76, AUC=0.78, *d*=0.60 |
| | | | | Self vs Control | acc.=0.71(+/-0.11), *P*=0.1435, sens.=0.71, spec.=0.71, AUC=0.70, *d*=0.39 |
| Validation | Study 5c | Bipolar patient | 18 | Self vs Other | acc.=0.78(+/-0.10), *P*=0.0309***, sens.=0.78, spec.=0.78, AUC=0.74, *d*=0.58 |
| | | | | Self vs Control | acc.=0.67(+/-0.11), *P*>0.20, sens.=0.67, spec.=0.67, AUC=0.71, *d*=0.35 |

***Other-Mentalizing Signature (MS-Other)***

| Signature | Dataset | Sample | n | Classification Task | Prediction Outcome |
|---|---|---|---|---|---|
| Training | Study 1 | Healthy adult | 21 | Other vs Self | acc.=1.00(+/-0.00), *P*<0.0001***, sens.=1.00, spec.=1.00, AUC=1.00, *d*=2.36 |
| | | | | Other vs Control | acc.=1.00(+/-0.00), *P*<0.0001***, sens.=1.00, spec.=1.00, AUC=1.00, *d*=2.53 |
| Validation | Study 2 | Adolescent | 44 | Other vs Self | acc.=0.70(+/-0.07), *P*=0.0096***, sens.=0.70, spec.=0.70, AUC=0.83, *d*=0.90 |
| | | | | Other vs Control | acc.=0.95(+/-0.03), *P*<0.0001***, sens.=0.95, spec.=0.95, AUC=0.97, *d*=0.99 |
| Validation | Study 3 | Adolescent | 61 | Other vs Self | acc.=0.75(+/-0.06), *P*<0.0001***, sens.=0.75, spec.=0.75, AUC=0.80, *d*=0.81 |
| | | | | Other vs Control | acc.=0.38(+/-0.06), *P*>0.20, sens.=0.38, spec.=0.38, AUC=0.39, *d*=0.39 |
| Validation | Study 4a | Healthy adult | 33 | Other vs Self | acc.=0.94(+/-0.04), *P*<0.0001***, sens.=0.94, spec.=0.94, AUC=0.99, *d*=2.15 |
| | | | | Other vs Control | acc.=0.88(+/-0.06), *P*<0.0001***, sens.=0.88, spec.=0.88, AUC=0.95, *d*=1.55 |

| | | | | | |
|---|---|---|---|---|---|
| Validation | Study 4b | Schizophrenia patient | 23 | Other vs Self | acc.=0.70(+/-0.10), *P*=0.0931, sens.=0.70, spec.=0.70, AUC=0.81, *d*=0.83 |
| | | | | Other vs Control | acc.=0.83(+/-0.08), *P*=0.0026***, sens.=0.83, spec.=0.83, AUC=0.90, *d*=1.21 |
| Validation | Study 5a | Healthy adult | 15 | Other vs Self | acc.=0.87(+/-0.09), *P*=0.0074***, sens.=0.87, spec.=0.87, AUC=0.97, *d*=1.65 |
| | | | | Other vs Control | acc.=1.00(+/-0.00), *P*<0.0001***, sens.=1.00, spec.=1.00, AUC=1.00, *d*=2.73 |
| Validation | Study 5b | Schizophrenia patient | 17 | Other vs Self | acc.=0.76(+/-0.10), *P*=0.0490***, sens.=0.76, spec.=0.76, AUC=0.93 , *d*=1.34 |
| | | | | Other vs Control | acc.=0.94(+/-0.06), *P*=0.0003***, sens.=0.94, spec.=0.94, AUC=0.99, *d*=1.89 |
| Validation | Study 5c | Bipolar patient | 18 | Other vs Self | acc.=0.83(+/-0.09), *P*=0.0075***, sens.=0.83, spec.=0.83, AUC=0.86, *d*=1.02 |
| | | | | Other vs Control | acc.=0.94(+/-0.05), *P*=0.0002***, sens.=0.94, spec.=0.94, AUC=0.96, *d*=1.79 |

***Mentalizing Signature (MS)***

| Signature | Dataset | Sample | n | Classification Task | Prediction Outcome |
|---|---|---|---|---|---|
| Training | Study 1 | Healthy adult | 21 | Self vs Control | acc.=1.00(+/-0.00), *P*<0.0001***, sens.=1.00, spec.=1.00, AUC=1.00, *d*=5.12 |
| | | | | Other vs Control | acc.=1.00(+/-0.00), *P*<0.0001***, sens.=1.00, spec.=1.00, AUC=1.00, *d*=4.72 |
| Validation | Study 2 | Adolescent | 44 | Self vs Control | acc.=1.00(+/-0.00), *P*<0.0001***, sens.=1.00, spec.=1.00, AUC=1.00, *d*=4.84 |
| | | | | Other vs Control | acc.=1.00(+/-0.00), *P*<0.0001***, sens.=1.00, spec.=1.00, AUC=1.00, *d*=4.99 |
| Validation | Study 3 | Adolescent | 61 | Self vs Control | acc.=1.00(+/-0.00), *P*<0.0001***, sens.=1.00, spec.=1.00, AUC=1.00, *d*=3.76 |
| | | | | Other vs Control | acc.=1.00(+/-0.00), *P*<0.0001***, sens.=1.00, spec.=1.00, AUC=1.00, *d*=3.98 |
| Validation | Study 4a | Healthy adult | 33 | Self vs Control | acc.=0.97(+/-0.03), *P*<0.0001***, sens.=0.97, spec.=0.97, AUC=1.00, *d*=2.13 |
| | | | | Other vs Control | acc.=0.97(+/-0.03), *P*<0.0001***, sens.=0.97, spec.=0.97, AUC=0.98, *d*=1.87 |
| Validation | Study 4b | Schizophrenia patient | 23 | Self vs Control | acc.=0.91(+/-0.06), *P*<0.0001***, sens.=0.91, spec.=0.91, AUC=0.99, *d*=1.61 |
| | | | | Other vs Control | acc.=0.96(+/-0.04), *P*<0.0001***, sens.=0.96, spec.=0.96, AUC=0.98, *d*=1.74 |
| Validation | Study 5a | Healthy adult | 15 | Self vs Control | acc.=1.00(+/-0.00), *P*<0.0001***, sens.=1.00, spec.=1.00, AUC=1.00, *d*=2.47 |
| | | | | Other vs Control | acc.=1.00(+/-0.00), *P*<0.0001***, sens.=1.00, spec.=1.00, AUC=1.00, *d*=3.36 |
| Validation | Study 5b | Schizophrenia patient | 17 | Self vs Control | acc.=1.00(+/-0.00), *P*<0.0001***, sens.=1.00, spec=1.00, AUC=1.00, *d*=1.95 |
| | | | | Other vs Control | acc.=0.94(+/-0.06), *P*=0.0003***, sens.=0.94, spec.=0.94, AUC=0.98, *d*=1.91 |
| Validation | Study 5c | Bipolar patient | 18 | Self vs Control | acc.=0.89(+/-0.07), *P*=0.0013***, sens.=0.89, spec.=0.89, AUC=0.97, *d*=1.61 |
| | | | | Other vs Control | acc.=0.94(+/-0.05), *P*=0.0002***, sens.=0.94, spec.=0.94, AUC=0.99, *d*=2.19 |

***Self-vs-Other Referential Signature (SvO-RS)***

| Signature | Dataset | Sample | n | Classification Task | Prediction Outcome |
|---|---|---|---|---|---|
| Training | Study 1 | Healthy adult | 21 | Self vs Other | acc.=1.00(+/-0.00), *P*<0.0001***, sens.=1.00, spec.=1.00, AUC=1.00, *d*=2.45 |
| Validation | Study 2 | Adolescent | 44 | Self vs Other | acc.=0.73(+/-0.07), *P*=0.0037***, sens.=0.73, spec.=0.73, AUC=0.85, *d*=1.04 |

| | | | | | |
|---|---|---|---|---|---|
| Validation | Study 3 | Adolescent | 61 | Self vs Other | acc.=0.75(+/-0.06), *P*<0.0001***, sens.=0.75, spec.=0.75, AUC=0.80, *d*=0.82 |
| Validation | Study 4a | Healthy adult | 33 | Self vs Other | acc.=0.91(+/-0.05), *P*<0.0001***, sens.=0.91, spec.=0.91, AUC=0.99, *d*=2.13 |
| Validation | Study 4b | Schizophrenia patient | 23 | Self vs Other | acc.=0.70(+/-0.10), *P*=0.0931, sens.=0.70, spec.=0.70, AUC=0.78, *d*=0.70 |
| Validation | Study 5a | Healthy adult | 15 | Self vs Other | acc.=0.80(+/-0.10), *P*=0.0352, sens.=0.80, spec.=0.80, AUC=0.93, *d*=1.37 |
| Validation | Study 5b | Schizophrenia patient | 17 | Self vs Other | acc.=0.76(+/-0.10), *P*=0.0490, sens.=0.76, spec.=0.76, AUC=0.93, *d*=1.32 |
| Validation | Study 5c | Bipolar patient | 18 | Self vs Other | acc.=0.83(+/-0.09), *P*=0.0075, sens.=0.83, spec.=0.83, AUC=0.83, *d*=0.89 |

*Note.* The predictions of signatures were assigned using paired observations with a forced-choice principle. acc. = accuracy; sens. = sensitivity; spec. = specificity, AUC = area under the curve. *d* refers to the estimated *Cohen's d* calculated as the mean difference of true and false paired predictions divided by the pooled standard deviation of differences (where difference = input_values[binary_class]- input_values[~binary_class]). Asterisks (***) mark the significant classification accuracies with $p < .05$.

### Supplementary Table 7

*Prediction results of the ROI classifiers across training and validation datasets*

| ***ROI 1: mPFC*** | | | |
|---|---|---|---|
| **Trained for** | **Task** | **Sample** | **Prediction Outcome** |
| Predicting Self-mentalizing | Self vs Other | Training (n=21) | acc.=0.95(+/-0.05), *P*<0.0001***, sens.=0.95, spec.=0.95, AUC=1.00, *d* =2.17 |
| | | Validation (n=211) | acc.=0.66(+/-0.03), *P*<0.0001***, sens.=0.66, spec.=0.66, AUC=0.69, *d*=0.37 |
| | Self vs Control | Training (n=21) | acc.=1.00(+/-0.00), *P*<0.0001***, sens.=1.00, spec.=1.00, AUC=1.00, *d*=2.29 |
| | | Validation (n=211) | acc.=0.79(+/-0.03), *P*<0.0001***, sens.=0.79, spec.=0.79, AUC=0.88, *d*=0.96 |
| Predicting Other-mentalizing | Other vs Self | Training (n=21) | acc.=1.00(+/-0.00), *P*<0.0001***, sens.=1.00, spec.=1.00, AUC=1.00, *d*=2.23 |
| | | Validation (n=211) | acc.=0.63(+/-0.03), *P*=0.0002***, sens.=0.63, spec.=0.63, AUC=0.66, *d*=0.36 |
| | Other vs Control | Training (n=21) | acc.=0.86(+/-0.08), *P*=0.0015***, sens.=0.86, spec.=0.86, AUC=0.97, *d*=1.68 |
| | | Validation (n=211) | acc.=0.78(+/-0.03), *P*<0.0001***, sens.=0.78, spec.=0.78, AUC=0.85, *d*=0.86 |
| Predicting mentalizing | Self vs Control | Training (n=21) | acc.=1.00(+/-0.00), *P*<0.0001***, sens.=1.00, spec.=1.00, AUC=1.00, *d*=5.63 |
| | | Validation (n=211) | acc.=0.97(+/-0.01), *P*<0.0001***, sens.=0.97, spec.=0.97, AUC=0.99, *d*=1,64 |
| | Other vs Control | Training (n=21) | acc.=1.00(+/-0.00), *P*<0.0001***, sens.=1.00, spec.=1.00, AUC=1.00, *d*=6.01 |
| | | Validation (n=211) | acc.=0.97(+/-0.01), *P*<0.0001***, sens.=0.97, spec.=0.97, AUC=0.99, *d*=1.69 |
| Differentiating „self" vs „other" conditions | Self vs Other | Training (n=21) | acc.=0.95(+/-0.05), *P*<0.0001***, sens.=0.95, spec.=0.95, AUC=1.00, *d*=2.10 |
| | | Validation (n=211) | acc.=0.68(+/-0.03), *P*<0.0001***, sens.=0.68, spec.=0.68, AUC=0.68, *d*=0.36 |
| ***ROI 2: Precuneus/PCC*** | | | |
| **Trained for** | **Task** | **Sample** | **Prediction Outcome** |
| Predicting Self-mentalizing | Self vs Other | Training (n=21) | acc.=0.81(+/-0.09), *P*=0.0072***, sens.=0.81, spec.=0.81, AUC=0.90, *d*=1.20 |
| | | Validation (n=211) | acc.=0.70(+/-0.03), *P*<0.0001***, sens.=0.70, spec.=0.70, AUC=0.74, *d*=0.57 |
| | Self vs Control | Training (n=21) | acc.=0.90(+/-0.06), *P*=0.0002***, sens.=0.90, spec.=0.90, AUC=0.96, *d*=1.52 |
| | | Validation (n=211) | acc.=0.68(+/-0.03), *P*<0.0001***, sens.=0.68, spec.=0.68, AUC=0.73, *d*=0.59 |
| Predicting Other-mentalizing | Other vs Self | Training (n=21) | acc.=1.00(+/-0.00), *P*<0.0001***, sens.=1.00, spec.=1.00, AUC=1.00, *d*=2.10 |
| | | Validation (n=211) | acc.=0.81(+/-0.03), *P*<0.0001***, sens.=0.81, spec.=0.81, AUC=0.84, *d*=0.85 |
| | Other vs Control | Training (n=21) | acc.=0.95(+/-0.05), *P*<0.0001***, sens.=0.95, spec.=0.95, AUC=1.00, *d*=2.72 |
| | | Validation (n=211) | acc.=0.91(+/-0.02), *P*<0.0001***, sens.=0.91, spec.=0.91, AUC=0.96, *d*=1.45 |
| Predicting mentalizing | Self vs Control | Training (n=21) | acc.=1.00(+/-0.00), *P*<0.0001***, sens.=1.00, spec.=1.00, AUC=1.00, *d*=3.11 |
| | | Validation (n=211) | acc.=0.91(+/-0.02), *P*<0.0001***, sens.=0.91, spec.=0.91, AUC=0.96, *d*=1.29 |
| | Other vs Control | Training (n=21) | acc.=1.00(+/-0.00), *P*<0.0001***, sens.=1.00, spec.=1.00, AUC=1.00, *d*=3.42 |

| | | | |
|---|---|---|---|
| | | Validation (n=211) | acc.=0.91(+/-0.02), *P*<0.0001***, sens.=0.91, spec.=0.91, AUC=0.97, *d*=1.40 |
| Differentiating „self" vs „other" conditions | Self vs Other | Training (n=21) | acc.=1.00(+/-0.00), *P*<0.0001***, sens.=1.00, spec.=1.00, AUC=1.00, *d*=2.07 |
| | | Validation (n=211) | acc.=0.75(+/-0.03), *P*<0.0001***, sens.=0.75, spec.=0.75, AUC=0.82, *d*=0.78 |

***ROI 3: TPJ right***

| Trained for | Task | Sample | Prediction Outcome |
|---|---|---|---|
| Predicting Self-mentalizing | Self vs Other | Training (n=21) | acc.=0.81(+/-0.09), *P*=0.0072***,sens.=0.81, spec.=0.81, AUC=0.84, *d*=0.69 |
| | | Validation (n=211) | acc.=0.47(+/-0.03), *P*>0.20, sens.=0.47, spec.=0.47, AUC=0.49, *d*=0.05 |
| | Self vs Control | Training (n=21) | acc.=0.76(+/-0.09), *P*=0.0266***, sens.=0.76, spec.=0.76, AUC=0.88, *d*=1.03 |
| | | Validation (n=211) | acc.=0.63(+/-0.03), *P*=0.0003***, sens.=0.63, spec.=0.63, AUC=0.69, *d*=0.45 |
| Predicting Other-mentalizing | Other vs Self | Training (n=21) | acc.=0.76(+/-0.09), *P*=0.0266***, sens.=0.76, spec.=0.76, AUC=0.86, *d*=0.90 |
| | | Validation (n=211) | acc.=0.69(+/-0.03), *P*<0.0001***, sens.=0.69, spec.=0.69, AUC=0.74, *d*=0.49 |
| | Other vs Control | Training (n=21) | acc.=0.81(+/-0.09), *P*=0.0072***, sens.=0.81, spec.=0.81, AUC=0.92, *d*=1.12 |
| | | Validation (n=211) | acc.=0.69(+/-0.03), *P*<0.0001***, sens.=0.69, spec.=0.69, AUC=0.74, *d*=0.60 |
| Predicting mentalizing | Self vs Control | Training (n=21) | acc.=1.00(+/-0.00), *P*<0.0001***, sens.=1.00, spec.=1.00, AUC=1.00, *d*=3.32 |
| | | Validation (n=211) | acc.=0.80(+/-0.03), *P*<0.0001***, sens.=0.80, spec.=0.80, AUC=0.90, *d*=1.07 |
| | Other vs Control | Training (n=21) | acc.=1.00(+/-0.00), *P*<0.0001***, sens.=1.00, spec.=1.00, AUC=1.00, *d*=2.55 |
| | | Validation (n=211) | acc.=0.86(+/-0.02), *P*<0.0001***, sens.=0.86, spec.=0.86, AUC=0.93, *d*=1.15 |
| Differentiating „self" vs „other" conditions | Self vs Other | Training (n=21) | acc.=0.76(+/-0.09), *P*=0.0266***, sens.=0.76, spec.=0.76, AUC=0.84, *d*=0.82 |
| | | Validation (n=211) | acc.=0.65(+/-0.03), *P*<0.0001***, sens.=0.65, spec.=0.65, AUC=0.71, *d*=0.45 |

***ROI 4: TPJ left***

| Trained for | Task | Sample | Prediction Outcome |
|---|---|---|---|
| Predicting Self-mentalizing | Self vs Other | Training (n=21) | acc.=0.57(+/-0.11), *P*>0.20, sens.=0.57, spec.=0.57, AUC=0.64, *d*=0.23 |
| | | Validation (n=211) | acc.=0.56(+/-0.03), *P*=0.0983, sens.=0.56, spec.=0.56, AUC=0.60, *d*=0.24 |
| | Self vs Control | Training (n=21) | acc.=0.81(+/-0.09), *P*=0.0072***, sens.=0.81, spec.=0.81, AUC=0.75, *d*=0.57 |
| | | Validation (n=211) | acc.=0.69(+/-0.03), *P*<0.0001***, sens.=0.69, spec.=0.69, AUC=0.80, *d*=0.74 |
| Predicting Other-mentalizing | Other vs Self | Training (n=21) | acc.=0.81(+/-0.09), *P*=0.0072***, sens.=0.81, spec.=0.81, AUC=0.87, *d*=0.95 |
| | | Validation (n=211) | acc.=0.71(+/-0.03), *P*<0.0001***, sens.=0.71, spec.=0.71, AUC=0.72, *d*=0.44 |
| | Other vs Control | Training (n=21) | acc.=0.90(+/-0.06), *P*=0.0002***, sens.=0.90, spec.=0.90, AUC=0.98, *d*=1.81 |
| | | Validation (n=211) | acc.=0.81(+/-0.03), *P*<0.0001***, sens.=0.81, spec.=0.81, AUC=0.90, *d*=1.07 |
| Predicting mentalizing | Self vs Control | Training (n=21) | acc.=1.00(+/-0.00), *P*<0.0001***, sens.=1.00, spec.=1.00, AUC=1.00, *d*=3.04 |

| | | | |
|---|---|---|---|
| | | Validation (n=211) | acc.=0.89(+/-0.02), *P*<0.0001***, sens.=0.89, spec.=0.89, AUC=0.96, *d*=1.19 |
| | Other vs Control | Training (n=21) | acc.=1.00(+/-0.00), *P*<0.0001***, sens.=1.00, spec.=1.00, AUC=1.00, *d*=2.70 |
| | | Validation (n=211) | acc.=0.92(+/-0.02), *P*<0.0001***, sens.=0.92, spec.=0.92, AUC=0.98, *d*=1.23 |
| Differentiating „self“ vs „other“ conditions | Self vs Other | Training (n=21) | acc.=0.86(+/-0.08), *P*=0.0015***, sens.=0.86, spec.=0.86, AUC=0.88, *d*=1.04 |
| | | Validation (n=211) | acc.=0.68(+/-0.03), *P*<0.0001***, sens.=0.68, spec.=0.68, AUC=0.72, *d*=0.45 |

***ROI 5: aMTG right***

| **Trained for** | **Task** | **Sample** | **Prediction Outcome** |
|---|---|---|---|
| Predicting Self-mentalizing | Self vs Other | Training (n=21) | acc.=0.67(+/-0.10), *P*=0.1893, sens.=0.67, spec.=0.67, AUC=0.74, *d*=0.51 |
| | | Validation (n=211) | acc.=0.64(+/-0.03), *P*<0.0001***, sens.=0.64, spec.=0.64, AUC=0.69, *d*=0.47 |
| | Self vs Control | Training (n=21) | acc.=0.67(+/-0.10), *P*=0.1893, sens.=0.67, spec.=0.67, AUC=0.75, *d*=0.59 |
| | | Validation (n=211) | acc.=0.41(+/-0.03), *P*>0.20, sens.=0.41, spec.=0.41, AUC=0.36, *d*=-0.37 |
| Predicting Other-mentalizing | Other vs Self | Training (n=21) | acc.=0.81(+/-0.09), *P*=0.0072***, sens.=0.81, spec.=0.81, AUC=0.86, *d*=1.11 |
| | | Validation (n=211) | acc.=0.66(+/-0.03), *P*<0.0001***, sens.=0.66, spec.=0.66, AUC=0.69, *d*=0.46 |
| | Other vs Control | Training (n=21) | acc.=0.81(+/-0.09), *P*=0.0072***, sens.=0.81, spec.=0.81, AUC=0.94, *d*=1.35 |
| | | Validation (n=211) | acc.=0.84(+/-0.03), *P*<0.0001***, sens.=0.84, spec.=0.84, AUC=0.92, *d*=1.13 |
| Predicting mentalizing | Self vs Control | Training (n=21) | acc.=0.76(+/-0.09), *P*=0.0266***, sens.=0.76, spec.=0.76, AUC=0.92, *d*=1.25 |
| | | Validation (n=211) | acc.=0.80(+/-0.03), *P*<0.0001***, sens.=0.80, spec.=0.80, AUC=0.87, *d*=1.00 |
| | Other vs Control | Training (n=21) | acc.=0.86(+/-0.08), *P*=0.0015***, sens.=0.86, spec.=0.86, AUC=0.95, *d*=1.39 |
| | | Validation (n=211) | acc.=0.83(+/-0.03), *P*<0.0001***, sens.=0.83, spec.=0.83, AUC=0.89, *d*=1.00 |
| Differentiating „self“ vs „other“ conditions | Self vs Other | Training (n=21) | acc.=0.81(+/-0.09), *P*=0.0072***, sens.=0.81, spec.=0.81, AUC=0.84, *d*=0.90 |
| | | Validation (n=211) | acc.=0.65(+/-0.03), *P*<0.0001***, sens.=0.65, spec.=0.65, AUC=0.71, *d*=0.51 |

***ROI 6: aMTG left***

| **Trained for** | **Task** | **Sample** | **Prediction Outcome** |
|---|---|---|---|
| Predicting Self-mentalizing | Self vs Other | Training (n=21) | acc.=0.95(+/-0.05), *P*<0.0001***, sens.=0.95, spec.=0.95, AUC=0.98, *d*=1.83 |
| | | Validation (n=211) | acc.=0.57(+/-0.03), *P*=0.0386***, sens.=0.57, spec.=0.57, AUC=0.63, *d*=0.35 |
| | Self vs Control | Training (n=21) | acc.=1.00(+/-0.00), *P*<0.0001***, sens.=1.00, spec.=1.00, AUC=1.00, *d*=2.53 |
| | | Validation (n=211) | acc.=0.75(+/-0.03), *P*<0.0001***, sens.=0.75, spec.=0.75, AUC=0.81, *d*=0.87 |
| Predicting Other-mentalizing | Other vs Self | Training (n=21) | acc.=0.86(+/-0.08), *P*=0.0015***, sens.=0.86, spec.=0.86, AUC=0.96, *d*=1.69 |
| | | Validation (n=211) | acc.=0.71(+/-0.03), *P*<0.0001***, sens.=0.71, spec.=0.71, AUC=0.80, *d*=0.81 |
| | Other vs Control | Training (n=21) | acc.=0.95(+/-0.05), *P*<0.0001***, sens.=0.95, spec.=0.95, AUC=0.97, *d*=1.71 |
| | | Validation (n=211) | acc.=0.73(+/-0.03), *P*<0.0001***, sens.=0.73, |

| | | | |
|---|---|---|---|
| | | | spec.=0.73, AUC=0.80, *d*=0.74 |
| Predicting mentalizing | Self vs Control | Training (n=21) | acc.=1.00(+/-0.00), *P*<0.0001***, sens.=1.00, spec.=1.00, AUC=1.00, *d*=4.43 |
| | | Validation (n=211) | acc.=0.92(+/-0.02), *P*<0.0001***, sens.=0.92, spec.=0.92, AUC=0.97, *d*=1.30 |
| | Other vs Control | Training (n=21) | acc.=1.00(+/-0.00), *P*<0.0001***, sens.=1.00, spec.=1.00, AUC=1.00, *d*=3.49 |
| | | Validation (n=211) | acc.=0.94(+/-0.02), *P*<0.0001***, sens.=0.94, spec.=0.94, AUC=0.98, *d*=1.47 |
| Differentiating „self" vs „other" conditions | Self vs Other | Training (n=21) | acc.=0.95(+/-0.05), *P*<0.0001***, sens.=0.95, spec.=0.95, AUC=0.99, *d*=1.91 |
| | | Validation (n=211) | acc.=0.64(+/-0.03), *P*<0.0001***, sens.=0.64, spec.=0.64, AUC=0.69, *d*=0.49 |

***ROI 7: SMA left***

| **Trained for** | **Task** | **Sample** | **Prediction Outcome** |
|---|---|---|---|
| Predicting Self-mentalizing | Self vs Other | Training (n=21) | acc.=0.67(+/-0.10), *P*=0.1893, sens.=0.67, spec.=0.67, AUC=0.70, *d*=0.23 |
| | | Validation (n=211) | acc.=0.52(+/-0.03), *P*>0.20, sens.=0.52, spec.=0.52, AUC=0.51, *d*=-0.02 |
| | Self vs Control | Training (n=21) | acc.=1.00(+/-0.00), *P*<0.0001***, sens.=1.00, spec.=1.00, AUC=1.00, *d*=2.44 |
| | | Validation (n=211) | acc.=0.48(+/-0.03), *P*>0.20, sens.=0.48, spec.=0.48, AUC=0.53, *d*=0.12 |
| Predicting Other-mentalizing | Other vs Self | Training (n=21) | acc.=0.43(+/-0.11), *P*>0.20, sens.=0.43, spec.=0.43, AUC=0.51, *d*=-0.02 |
| | | Validation (n=211) | acc.=0.50(+/-0.03), *P*>0.20, sens.=0.50, spec.=0.50, AUC=0.51, *d*=-0.08 |
| | Other vs Control | Training (n=21) | acc.=0.62(+/-0.11), *P*>0.20, sens.=0.62, spec.=0.62, AUC=0.81, *d*=0.87 |
| | | Validation (n=211) | acc.=0.71(+/-0.03), *P*<0.0001***, sens.=0.71, spec.=0.71, AUC=0.75, *d*=0.60 |
| Predicting mentalizing | Self vs Control | Training (n=21) | acc.=1.00(+/-0.00), *P*<0.0001***, sens.=1.00, spec.=1.00, AUC=1.00, *d*=4.30 |
| | | Validation (n=211) | acc.=0.85(+/-0.02), *P*<0.0001***, sens.=0.85, spec.=0.85, AUC=0.94, *d*=1.20 |
| | Other vs Control | Training (n=21) | acc.=0.95(+/-0.05), *P*<0.0001***, sens.=0.95, spec.=0.95, AUC=1.00, *d*=3.05 |
| | | Validation (n=211) | acc.=0.86(+/-0.02), *P*<0.0001***, sens.=0.86, spec.=0.86, AUC=0.93, *d*=1.20 |
| Differentiating „self" vs „other" conditions | Self vs Other | Training (n=21) | acc.=0.62(+/-0.11), *P*>0.20, sens.=0.62, spec.=0.62, AUC=0.69, *d*=0.20 |
| | | Validation (n=211) | acc.=0.50(+/-0.03), *P*>0.20, sens.=0.50, spec.=0.50, AUC=0.50, *d*=-0.03 |

***ROI 8: Cerebellum bilateral***

| **Trained for** | **Task** | **Sample** | **Prediction Outcome** |
|---|---|---|---|
| Predicting Self-mentalizing | Self vs Other | Training (n=21) | acc.=0.62(+/-0.11), *P*>0.20, sens.=0.62, spec.=0.62, AUC=0.65, *d*=0.29 |
| | | Validation (n=211) | acc.=0.52(+/-0.03), *P*>0.20, sens.=0.52, spec=0.52, AUC=0.56, *d*=0.10 |
| | Self vs Control | Training (n=21) | acc.=0.86(+/-0.08), *P*=0.0015***, sens.=0.86, spec.=0.86, AUC=0.94, *d*=1.48 |
| | | Validation (n=211) | acc.=0.67(+/-0.03), *P*<0.0001***, sens.=0.67, spec.=0.67, AUC=0.71, *d*=0.39 |
| Predicting Other-mentalizing | Other vs Self | Training (n=21) | acc.=0.48(+/-0.11), *P*>0.20, sens.=0.48, spec.=0.48, AUC=0.54, *d*=0.06 |
| | | Validation (n=211) | acc.=0.54(+/-0.03), *P*>0.20, sens.=0.54, spec.=0.54, AUC=0.55, *d*=0.05 |

| | | | |
|---|---|---|---|
| | Other vs Control | Training (n=21) | acc.=0.76(+/-0.09), *P*=0.0266***, sens.=0.76, spec.=0.76, AUC=0.85, *d*=0.81 |
| | | Validation (n=211) | acc.=0.45(+/-0.03), *P*>0.20, sens.=0.45, spec.=0.45, AUC=0.46, *d*=-0.10 |
| Predicting mentalizing | Self vs Control | Training (n=21) | acc.=1.00(+/-0.00), *P*<0.0001***, sens.=1.00, spec.=1.00, AUC=1.00, *d*=2.70 |
| | | Validation (n=211) | acc.=0.69(+/-0.03), *P*<0.0001***, sens.=0.69, spec.=0.69, AUC=0.76, *d*=0.61 |
| | Other vs Control | Training (n=21) | acc.=0.95(+/-0.05), *P*<0.0001***, sens.=0.95, spec.=0.95, AUC=1.00, *d*=2.61 |
| | | Validation (n=211) | acc.=0.73(+/-0.03), *P*<0.0001***, sens.=0.73, spec.=0.73, AUC=0.77, *d*=0.68 |
| Differentiating „self“ vs „other“ conditions | Self vs Other | Training (n=21) | acc.=0.62(+/-0.11), *P*>0.20, sens.=0.62, spec.=0.62, AUC=0.64, *d*=0.26 |
| | | Validation (n=211) | acc.=0.54(+/-0.03), *P*>0.20, sens.=0.54, spec.=0.54, AUC=0.56, *d*=0.07 |

***ROI 9: Cerebellum right***

| Trained for | Task | Sample | Prediction Outcome |
|---|---|---|---|
| Predicting Self-mentalizing | Self vs Other | Training (n=21) | acc.=0.48(+/-0.11), *P*>0.20, sens.=0.48, spec.=0.48, AUC=0.46, *d*=-0.31 |
| | | Validation (n=211) | acc.=0.40(+/-0.03), *P*>0.20, sens.=0.40, spec.=0.40, AUC=0.37, *d*=-0.34 |
| | Self vs Control | Training (n=21) | acc.=0.76(+/-0.09), *P*=0.0266***, sens.=0.76, spec.=0.76, AUC=0.84, *d*=0.95 |
| | | Validation (n=211) | acc.=0.73(+/-0.03), *P*<0.0001***, sens.=0.73, spec.=0.73, AUC=0.81, *d*=0.79 |
| Predicting Other-mentalizing | Other vs Self | Training (n=21) | acc.=0.67(+/-0.10), *P*=0.1893, sens.=0.67, spec.=0.67, AUC=0.62, *d*=0.21 |
| | | Validation (n=211) | acc.=0.49(+/-0.03), *P*>0.20, sens.=0.49, spec.=0.49, AUC=0.47, *d*=-0.10 |
| | Other vs Control | Training (n=21) | acc.=0.90(+/-0.06), *P*=0.0002***, sens.=0.90, spec.=0.90, AUC=0.95, *d*=1.31 |
| | | Validation (n=211) | acc.=0.60(+/-0.03), *P*=0.0058***, sens.=0.60, spec.=0.60, AUC=0.64, *d*=0.38 |
| Predicting mentalizing | Self vs Control | Training (n=21) | acc.=1.00(+/-0.00), *P*<0.0001***, sens.=1.00, spec.=1.00, AUC=1.00, *d*=3.52 |
| | | Validation (n=211) | acc.=0.88(+/-0.02), *P*<0.0001***, sens.=0.88, spec.=0.88, AUC=0.96, *d*=1.32 |
| | Other vs Control | Training (n=21) | acc.=1.00(+/-0.00), *P*<0.0001***, sens.=1.00, spec.=1.00, AUC=1.00, *d*=3.61 |
| | | Validation (n=211) | acc.=0.88(+/-0.02), *P*<0.0001***, sens.=0.88, spec.=0.88, AUC=0.95, *d*=1.36 |
| Differentiating „self“ vs „other“ conditions | Self vs Other | Training (n=21) | acc.=0.57(+/-0.11), *P*>0.20, sens.=0.57, spec.=0.57, AUC=0.51, *d*=-0.14 |
| | | Validation (n=211) | acc.=0.43(+/-0.03), *P*>0.20, sens.=0.43, spec.=0.43, AUC=0.41, *d*=-0.24 |

***ROI 10: Cerebellum left***

| Trained for | Task | Sample | Prediction Outcome |
|---|---|---|---|
| Predicting Self-mentalizing | Self vs Other | Training (n=21) | acc.=0.29(+/-0.10), *P*>0.20, sens.=0.29, spec.=0.29, AUC=0.45, *d*=-0.05 |
| | | Validation (n=211) | acc.=0.49(+/-0.03), *P*>0.20, sens.=0.49, spec.=0.49, AUC=0.50, *d*=-0.02 |
| | Self vs Control | Training (n=21) | acc.=0.76(+/-0.09), *P*=0.0266***, sens.=0.76, spec.=0.76, AUC=0.90, *d*=0.94 |
| | | Validation (n=211) | acc.=0.71(+/-0.03), *P*<0.0001***, sens.=0.71, spec.=0.71, AUC=0.78, *d*=0.73 |
| Predicting | Other vs Self | Training (n=21) | acc.=0.62(+/-0.11), *P*>0.20, sens.=0.62, |

| | | | |
|---|---|---|---|
| Other-mentalizing | | | spec.=0.62, AUC=0.63, *d*=0.12 |
| | | Validation (n=211) | acc.=0.52(+/-0.03), *P*>0.20, sens.=0.52, spec.=0.52, AUC=0.50, *d*=-0.02 |
| | Other vs Control | Training (n=21) | acc.=0.81(+/-0.09), *P*=0.0072***, sens.=0.81, spec.=0.81, AUC=0.88, *d*=1.09 |
| | | Validation (n=211) | acc.=0.62(+/-0.03), *P*=0.0006***, sens.=0.62, spec.=0.62, AUC=0.64, *d*=0.35 |
| Predicting mentalizing | Self vs Control | Training (n=21) | acc.=0.90(+/-0.06), *P*=0.0002***, sens.=0.90, spec.=0.90, AUC=0.99, *d*=2.26 |
| | | Validation (n=211) | acc.=0.85(+/-0.02), *P*<0.0001***, sens.=0.85, spec.=0.85, AUC=0.93, *d*=1.12 |
| | Other vs Control | Training (n=21) | acc.=0.90(+/-0.06), *P*=0.0002***, sens.=0.90, spec.=0.90, AUC=0.99, *d*=2.10 |
| | | Validation (n=211) | acc.=0.86(+/-0.02), *P*<0.0001***, sens.=0.86, spec.=0.86, AUC=0.93, *d*=1.08 |
| Differentiating „self“ vs „other“ conditions | Self vs Other | Training (n=21) | acc.=0.48(+/-0.11), *P*>0.20, sens.=0.48, spec.=0.48, AUC=0.58, *d*=0.11 |
| | | Validation (n=211) | acc.=0.48(+/-0.03), *P*>0.20, sens.=0.48, spec.=0.48, AUC=0.49, *d*=-0.04 |

*Note.* Each ROI was trained for four classification tasks and tested in validation datasets. acc. = accuracy; sens. = sensitivity, spec. = specificity, AUC = area under the curve. *d* refers to the estimated *Cohen's d* calculated as the mean difference of true and false paired predictions divided by the pooled standard deviation of differences (where difference = input_values[binary_class]- input_values[~binary_class]). Asterisks (***) mark the significant classification accuracies with $p < .05$.
